# A Goldilocks zone of DNA flexibility defines stable yet plastic nucleosomes, tuned by histone chemistry

**DOI:** 10.64898/2026.02.16.706184

**Authors:** Jose Ignacio Perez-Lopez, M. Julia Maristany, Stephen E. Farr, Jan Huertas, Rosana Collepardo-Guevara

## Abstract

Nucleosomes regulate DNA accessibility through partial DNA unwrapping driven by thermal fluctuations or forces exerted by molecular motors. Despite substantial chemical and DNA sequence diversity in vivo, much of our mechanistic understanding of nucleosome unwrapping is derived from canonical constructs. Here, we use a near-atomistic coarse-grained model to quantify the energetics and mechanisms of force-induced nucleosome unwrapping across over 40 nucleosomes spanning genomic and synthetic DNA sequences, histone variants, and post-translational modifications. We find that the force barriers to nucleosome unwrapping are shaped by histone composition and DNA flexibility, and arise from the formation of topologically protected, partially unwrapped intermediates. Strong nucleosome-positioning and genomic sequences consistently fall within a ‘Goldilocks zone’ of intermediate DNA flexibility, which enables the formation of thermodynamically stable yet mechanically plastic nucleosomes that undergo controlled deformation under applied force. Limited mechanical variation within this zone results in modest DNA sequence-dependent effects on nucleosome deformability. In contrast, histone variants and post-translational modifications modulate nucleosome stability and plasticity in a strongly non-additive manner, with predominant contributions from H3 and H2A core arginines and H3 and H2B tail lysines. Together, our results provide a physical framework linking DNA sequence, histone composition, and nucleosome geometry to nucleosome stability, plasticity and DNA accessibility.

## Introduction

DNA transcription, replication, and repair require transient access to the genetic material. In eukaryotic cells, however, this access is intrinsically constrained because DNA is not found free in solution but is compacted into a DNA–protein polymer known as chromatin, whose basic repeating unit is the nucleosome^1^. Interactions among nucleosomes and with other biomolecules promote chromatin compaction^2^. In addition, nucleosomes restrict DNA accessibility directly by wrapping DNA around a histone core, thereby occluding binding sites and limiting access by DNA-binding proteins. Controlling DNA access, therefore, relies on molecular machinery capable of repositioning nucleosomes or dynamically unwrapping and rewrapping nucleosomal DNA on demand^3^. Understanding what makes a nucleosome easy or hard to unwrap—and how its chemical composition modulates this response—is fundamental to deciphering gene function at the molecular level.

Structurally, a nucleosome consists of ∼147 base pairs (bp) of DNA wrapped in approximately 1.7 turns around a histone octamer composed of two copies each of histones H2A, H2B, H3, and H4^1,4^. The wrapped DNA is conventionally divided into two superhelical turns: the outer (first) turn, corresponding to the DNA distal from the dyad axis, and the inner (second) turn, corresponding to the DNA proximal to the dyad^5^. These regions differ in accessibility, with the outer turn being more readily unwrapped than the inner turn.

Ten intrinsically disordered, positively charged disordered protein regions, known as the histone tails, extend from the nucleosome core^1^. These include one N-terminal tail from each of the eight histone subunits, together with a C-terminal tail from each of the two H2A proteins. These flexible tails establish associative electrostatic interactions with DNA and with other chromatin components, contributing to nucleosome stability, higher-order chromatin structure^6–9^ and chromatin phase separation^10,11^.

Far from being rigid particles, nucleosomes undergo constant structural fluctuations^12,13^. A prominent example is nucleosome breathing: the transient partial unwrapping and rewrapping of DNA driven by thermal fluctuations^12,13^. Breathing occurs even in canonical nucleosomes and is enhanced at physiological salt concentrations^14,15^. By transiently exposing otherwise buried DNA motifs, nucleosome breathing is thought to facilitate access to transcription factors and other chromatin-associated proteins^16–19^. Superimposed on this intrinsic dynamics, nucleosomes are chemically heterogeneous and dynamically regulated through the incorporation of histone variants^20,21^ and post-translational modifications (PTMs)^22^. These chemical modifications can reshape the DNA–histone interaction interface and, thus, modulate the local stability, structure, and breathing dynamics of individual nucleosomes^23–26^. Hence, by modulating nucleosome dynamics and its chemical composition, these modifications impact the strength of nucleosome–nucleosome interactions^6,7,9,27–29^, the recruitment of chromatin-associated proteins, the structure of chromatin^6^, and its phase behaviour^10,30^.

Despite the importance of the chemical diversity of nucleosomes, our mechanistic understanding of how nucleosome stability is modulated in a DNA sequence- and chemical-composition-dependent manner remains incomplete. Here, we address this gap by combining equilibrium umbrella sampling molecular dynamics simulations of nucleosomes with our near-atomistic coarse-grained model of chromatin^30^. Using this approach, we quantify the energetics of nucleosome unwrapping under applied force across more than 40 different nucleosomes, spanning genomic and synthetic DNA sequences, histone variants, and acetylation patterns. This corresponds to an aggregate sampling time of over 700 *µ*s of near-atomistic nucleosome simulations. Our approach yields free-energy landscapes, force-vs-extension pulling curves, and near-atomistic structural snapshots of partially unwrapped states across the unwrapping process.

We show that the force barriers observed during nucleosome unwrapping arise from partially unwrapped ‘topologically protected’ intermediates in which the configuration of the linker DNA and the orientation of the nucleosome relative to the pulling direction prevent further extension without a global reorientation. The stability of these intermediates governs both the height of the free-energy barriers and the range in extension over which they persist, thereby determining the total reversible work required to convert the wrapped nucleosome into the fully unwrapped state. This stability is governed by a balance between sequence-dependent DNA mechanics, nucleosome orientation, and both associative electrostatic histone–DNA and repulsive DNA–DNA interactions. Chemical modifications to histones, including acetylation and histone variants, strongly modulate this balance in a highly non-additive and context-dependent manner, with specific marks and variants exerting effects that cannot be explained by simple charge neutralisation.

In parallel, we find that DNA sequence modulates nucleosome mechanics by tuning both the energetic cost of wrapping and the mechanical response of the partially unwrapped, topologically protected states. This defines a ‘Goldilocks zone’ of intermediate DNA flexibility, in which DNA is sufficiently flexible to support thermodynamically stable nucleosome assembly, while retaining enough stiffness to permit controlled deformation and unwrapping under applied force. Notably, both strong positioning sequences and genomic nucleosomal sequences consistently fall within this intermediate zone. Together, our results establish nucleosome mechanics as an emergent property arising from the coupled interplay between DNA elasticity, histone chemistry, and nucleosome geometry.

Overall, our work provides a physical framework for understanding the molecular determinants of nucleosome stability and plasticity, revealing how the physicochemical diversity of DNA sequences and histone chemistries encodes a broad range of mechanically distinct and regulable nucleosome behaviours.

## Results

### Nucleosome unwrapping with coarse-grained simulations at near-atomistic resolution

To investigate how the chemical composition of the nucleosome particle influences its stability, we quantify the change in free energy associated with nucleosome unwrapping under an applied force using umbrella sampling coarse-grained simulations. These simulations employ our near-atomistic, chemically specific, coarse-grained chromatin model^30^, which represents proteins at residue resolution and DNA at the base-pair level. We have previously shown^30^ that this model yields nucleosomes that undergo force-induced unwrapping in agreement with various single-molecule experiments^5,31–33^.

During in vitro force spectroscopy experiments, nucleosomes are pulled at near-equilibrium speeds (e.g., 0.1 mm/s for magnetic tweezers^33^), which are too slow to be reproduced directly by brute-force MD simulations. Capturing the equilibrium mechanism of DNA unwrapping would require millisecond-scale trajectories, which are infeasible with current computing power. To overcome this limitation, we perform equilibrium umbrella sampling simulations. Combined with the weighted histogram analysis method (WHAM)^34^, this approach allows us to calculate the Landau free energy profile, or Potential of Mean Force (PMF), of nucleosome unwrapping along a reaction coordinate.

We selected the DNA extension (i.e., the end-to-end distance between the first and last DNA base pairs) as the reaction coordinate, as it effectively captures the progression of nucleosome unwrapping^30,35^. We obtained force–extension (‘pulling’) curves by numerically differentiating the PMF with respect to this coordinate. Throughout this work, we report the external pulling force; i.e., the instantaneous mechanical force required to induce further unwrapping, which corresponds to the positive derivative of the PMF with respect to the extension. This definition has the opposite sign to the force exerted by the nucleosome itself to pull the DNA back in. In addition, we quantify the frequency of histone–DNA contacts along the DNA and analyse the reorientation angle, which is defined as the angle *β* between a vector normal to the nucleosome face and the pulling direction, as defined previously^36^. The reorientation angle serves as a geometric indicator of how nucleosome orientation, and in turn, DNA topology, modulate the mechanical efficiency of DNA unwrapping under tension.

We first applied this approach to a well-characterised reference nucleosome assembled from canonical histones taken from the 1KX5 crystal structure^37^. The associated DNA comprised a 211-bp construct: the 147-bp nucleosomal DNA sequence flanked symmetrically by two 32-bp linker DNA arms. These linker sequences were taken from the 1ZBB tetranucleosome crystal structure^38^. The quasi-palindromic 1KX5 DNA sequence^37^ makes this nucleosome an ideal benchmark for validating our model before testing histone mutations or sequence variations. For clarity, we centre the dyad at base pair 0 and define the outer turn as DNA spanning base pairs ±73 to ±36, and the inner turn as base pairs ±35 to 0. We refer to base pairs –105 to 0 as the left arm and 0 to +105 as the right arm, with the outermost regions (base pairs ±105 to ±74) corresponding to the flanking DNA linkers.

Analysis of the PMF (Fig. 1A) and its numerical derivative (Fig. 1B), which represents the force exerted on the system, reveals four distinct stages of nucleosome unwrapping, each marked by an approximately linear segment in the PMF with a distinct effective slope and a corresponding shift in the force.

**Figure 1.**
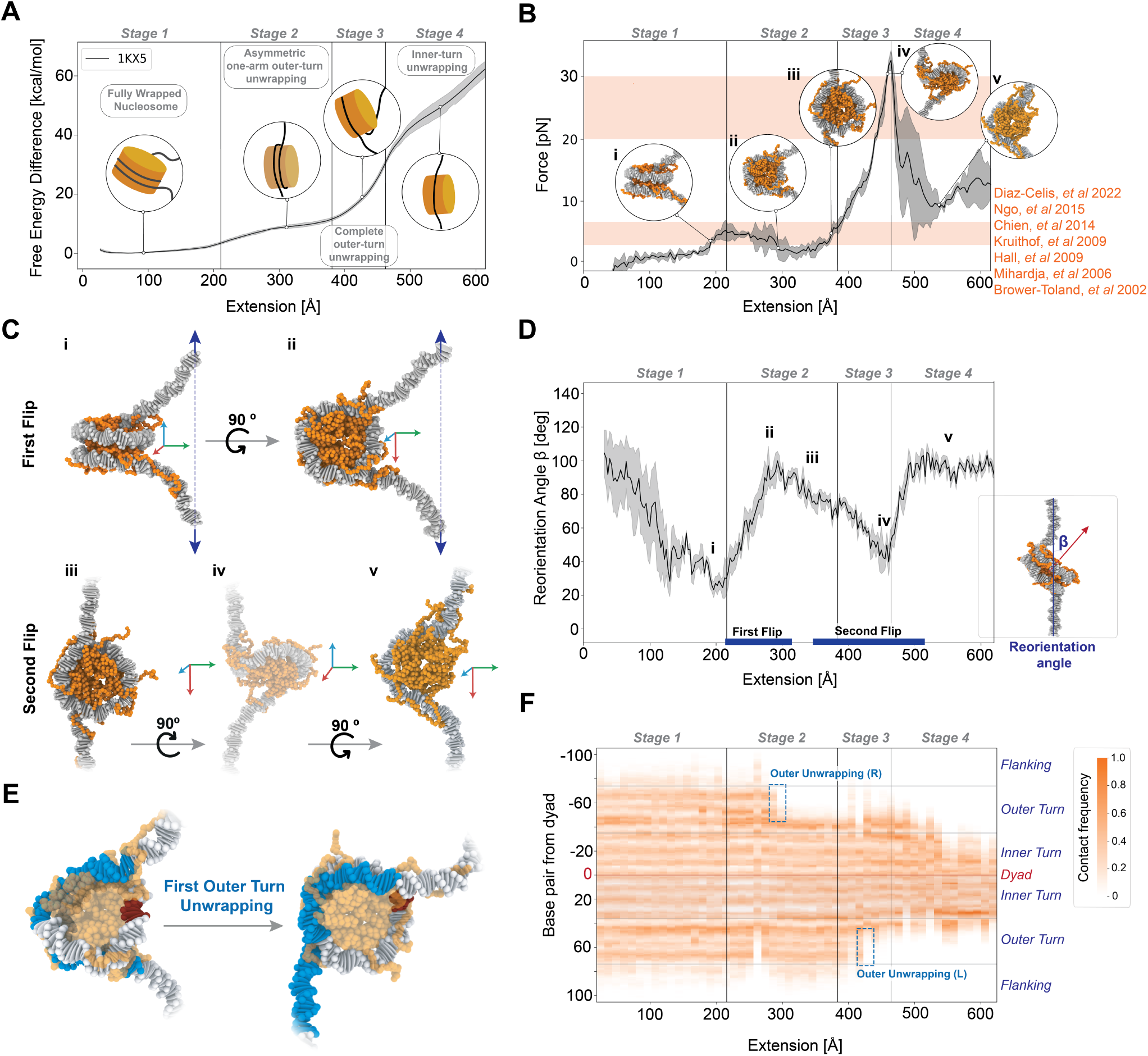
Nucleosome unwrapping under tension follows a multi-stage mechanism in which topologically protected partially-unwrapped intermediates control force barriers. **A.** Free-energy difference as a function of end-to-end extension for a 1KX5 nucleosome containing 211 base pairs. Four distinct unwrapping stages are indicated: **Stage 1,** fully wrapped nucleosome; **Stage 2,** asymmetric unwrapping of one arm of the outer DNA turn; **Stage 3,** unwrapping of the second arm of the outer DNA turn; and **Stage 4,** inner-turn unwrapping. **B.** Force–extension unwrapping curve obtained by taking the numerical derivative of panel A. The force maxima separating the unwrapping stages are indicated, together with representative experimental values reported previously^31,32,39–42,44^. **C.** Representative simulation snapshots illustrating two reorientation (‘flip’) transitions during force-induced unwrapping. Panels (i)–(ii) show the first reorientation, occurring during Stage 2, in which the nucleosome flips, aligning its face with the pulling direction, enabling the onset of outer-turn DNA release. Panels (iii)–(v) depict the second reorientation, occurring during Stage 4, which enables unwrapping of the inner DNA turn. Arrows indicate the reference frame and the direction of DNA extension; DNA is shown in grey and the histone octamer in orange. **D.** Reorientation angle *β*, defined as the angle between the normal vector to the plane of the nucleosome face and the pulling direction, plotted as a function of end-to-end extension. **E.** Structural snapshots illustrating the onset of asymmetric unwrapping of the first outer DNA turn. As tension increases, DNA detachment initiates stochastically from one of the two arms. Outer-turn DNA beads are highlighted in blue, the dyad in red, remaining DNA in grey, and the histone octamer in orange. **F.** Base-pair–resolved DNA–histone contact map as a function of end-to-end extension. The heatmap shows the normalised contact frequency between the histone core and DNA; flanking DNA, outer-turn, inner-turn, and dyad regions are annotated. The grey shadow in panels A, B and D represents the standard error.

During the first stage, the DNA extension increases markedly (by ∼200 Å) while both the force and the free energy change remain very small (Stage 1; Fig. 1A, B). In this regime, the measured force stays below 1 pN and the free energy variation is negligible, as the increase in extension primarily reflects the straightening of the flanking DNA arms. Here, the reorientation angle is initially large because the simulations are started from configurations in which the DNA ends are constrained to a short end-to-end distance of 25 Å and the nucleosomal superhelical axis is aligned with the pulling direction (Fig. 1D).

Stage 2 begins when the applied force reaches a first local maximum of approximately 6–7 pN, consistent with characteristic forces observed in pulling experiments^31–33,39–42^ (Fig. 1A, B, Supplementary Table 1, Supplementary Video 1). Analyses of nucleosome configurations (Fig. 1C) and of the reorientation angle (Fig. 1D) indicate that this force maximum originates from a topologically ‘protected’ configuration. In this protected configuration, the nucleosome remains fully wrapped, with its face, and thus the nucleosomal superhelical path, oriented perpendicular to the direction of the applied force (Fig. 1D, state i). This orientation corresponds to a minimum in the reorientation angle as the vector normal to the nucleosome face is aligned with the pulling direction. This intermediate nucleosome state is mechanically unfavourable for initiating DNA release, resulting in a transient build-up of force.

For unwrapping to proceed, the nucleosome must undergo a reorientation (‘flipping’) transition to a state in which the nucleosome face and the nucleosomal DNA path become approximately parallel to the pulling direction (Fig. 1C, transition from state i to ii), corresponding to a maximum in the reorientation angle (Fig. 1D, state ii). This structural transition aligns the applied force optimally to drive DNA release. Strikingly, once this reorientation has occurred, unwrapping of the first few (∼10 bp) base pairs in the entry/exit region—the region of the DNA usually involved in nucleosome breathing—proceeds at low forces, resulting in only a modest increase in free energy as histone–DNA contacts begin to break.

Following this reorientation, Stage 2 is characterised by asymmetric unwrapping of the outer DNA turn, in which only one linker arm undergoes loss of histone–DNA contacts and detaches while the other remains bound (Fig. 1E,F; Outer Unwrapping (R)). Owing to the near-palindromic nature of the 1KX5 DNA sequence, the initiating arm (left or right) is selected stochastically. Across 100 independent simulation replicates, initiation occurred from the right arm in 47 cases, from the left in 49 cases, and was simultaneous in 4 cases (Fig. S2). Single-molecule experiments with the Widom 601 sequence have also reported asymmetric unwrapping. However, given the asymmetric nature of that sequence, those experiments observed a consistent preference for one arm^31,40,43^. Stage 3 is characterised by a steep rise in both free energy and force (Fig. 1A, B), corresponding to unwrapping of the second outer-turn arm (Fig. 1F; Outer Unwrapping (L)).

After full outer-turn release, the force required for further DNA unwrapping increases sharply, reaching a second and more pronounced maximum exceeding 30 pN, marking the onset of Stage 4 (Fig. 1A,B). This peak coincides with the formation of a second topologically protected intermediate, in which the applied force is again poorly aligned with the unwrapping direction and acts to tighten the DNA against the histone core, leading to a build-up of force (Fig. 1C, state iii). Importantly, unlike the first protected state, which can be bypassed at moderate force through an early flipping of the nucleosome, this second protected configuration constitutes a substantially larger mechanical barrier. Once the outer DNA turn has fully detached, the loss of electrostatic repulsion between adjacent DNA turns stabilises the wrapping of the remaining inner turn. Eventually, at high forces, the nucleosome flips (Fig. 1C, states iv and v, Supplementary Video 1), as indicated by reorientation angles close to 100^◦^ (Fig. 1D, state v). Following this flipping, unwrapping of the inner turn proceeds readily and histone–DNA contacts are progressively lost.

Our pulling simulations are quantitatively consistent with force values reported across a range of experimental and theoretical studies, which consistently show that the inner DNA turn resists unwrapping more strongly than the outer turn^31–33,36,39–41,44^ (Supplementary Table 1). Optical tweezers and force–FRET measurements typically report two characteristic force regimes: a low-force regime of ∼ 2–5 pN associated with outer-turn release, and an inner turn-related transition starting at *>*10 pN, which corresponds to irreversible unwrapping of the inner DNA gyre^31,40–42^. At larger extensions, where the DNA is tightened against the histone surface or when unfavourable nucleosome orientations persist, experiments have also reported higher-force rupture events in the 15–30 pN range^32,39,44^. In this context, the two maxima in our pulling curves, at approximately 6–7 pN and 35 pN, align with these experimentally observed force regimes.

Our observations are consistent with previous simulations and theoretical analyses^36,43,45–47^ proposing that the higher resistance of the inner DNA turn to unwrapping arises from a combination of factors, including geometric constraints, steric hindrance between DNA turns, and the reduction in DNA–DNA electrostatic repulsion once the outer turn has detached. Together, these effects have been suggested to stabilise a metastable intermediate in which a single DNA turn remains wrapped around the histone core, in agreement with the topologically protected partially unwrapped intermediate observed here. Experimentally, slower unwrapping rates for the inner turn have also been attributed to stronger histone–DNA contacts near the dyad, including cooperative hydrogen bonding^44^. In our model, all histone–DNA electrostatic interactions are treated equivalently, and cooperative hydrogen bonds are not explicitly represented. Yet we still observe a markedly higher second force peak, indicating that the mechanical bottleneck protecting the inner turn arises naturally from DNA mechanics, nucleosome orientation, and physical constraints. This supports the view that the second force maximum reflects the disruption of a topologically protected nucleosome intermediate^43^, rather than merely the breaking of site-specific histone–DNA contacts.

### Physicochemical regulation of nucleosome stability and plasticity at amino-acid resolution

Having established a detailed mechanism of force–induced nucleosome unwrapping, and having validated our model against experimental force–extension measurements, we next exploit its chemical specificity and sub-molecular resolution to identify the molecular interactions that govern nucleosome stability and its resistance to force-induced unwrapping.

The propensity of DNA to wrap around the histone core and form nucleosomes is understood to be dominated by two main factors: the intrinsic flexibility of the DNA and the associative interactions formed between DNA and histone residues. In this section, we focus on characterising the relative contributions of individual histone–DNA electrostatic interactions to nucleosome stability under applied force. In particular, we quantify these contributions by histone type (H2A, H2B, H3, H4), by structural region (globular domains versus histone tails), and by amino-acid identity (arginine versus lysine). To this end, we perform an energy-weighted contact analysis. For each arginine–DNA or lysine–DNA pair classified as ‘in contact’ (i.e., closer than a cut-off distance; see Methods), we calculate the corresponding electrostatic interaction energy according to the force field (Fig. 2A). We then plot the summed interaction energies as a function of the end-to-end DNA extension, decomposed according to the four nucleosome-unwrapping stages defined above.

**Figure 2.**
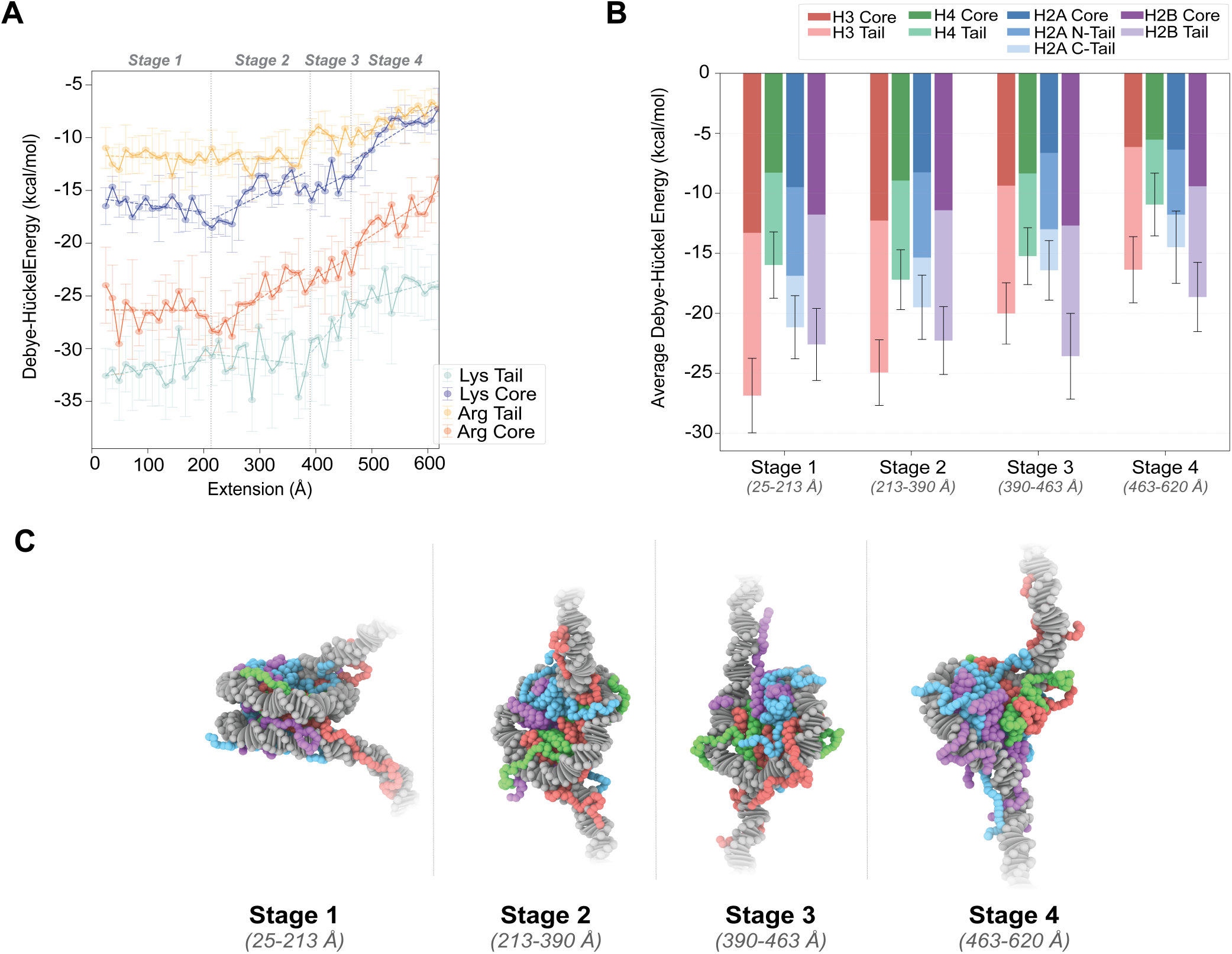
Histone core and tail residues play distinct, stage-dependent roles during force-induced nucleosome unwrapping. **A.** Debye–Hückel electrostatic interaction energy arising from arginine–DNA and lysine–DNA contacts as a function of the end-to-end extension. Contributions from histone cores and tails are shown separately. **B.** Average electrostatic interaction energy between DNA and individual histone components, grouped by unwrapping stage. Contributions from histone tails and cores are shown separately for each histone type. **C.** Representative configurations illustrating the stages of force-induced unwrapping, with histones highlighted (H3 red, H4 green, H2A blue, H2B purple). **Stage 1:** initial fully wrapped state. **Stage 2:** unwrapping-efficient nucleosome configuration enabling unwrapping of one of the outer-turn DNA arms. **Stage 3:** unwrapping of the second outer-turn DNA arms. **Stage 4:** unwrapping-efficient nucleosome configuration enabling inner-turn unwrapping.

In the fully wrapped nucleosome (Stage 1), stability is supported predominantly by strong associative interactions involving lysine residues within the flexible histone tails and arginine residues within the globular histone core, which contribute with comparable magnitude (Fig. 2A). In contrast, arginines in the tails and lysines in the core make more modest contributions (Fig. 2A). These trends indicate that lysines and arginines fulfill complementary roles in nucleosome stabilisation: core arginines act as relatively stable anchoring points, whereas tail lysines confer flexible and tuneable electrostatic screening.

The distinct roles of lysine and arginine residues in maintaining nucleosome stability are consistent with previous experiments and simulations^1,48,49^. In the 1KX5 nucleosome, the histone core contains a total of 82 arginine and 56 lysine residues, whereas the histone tails contain 22 arginine and 58 lysine residues. Therefore, the experimentally reported dominant contribution of arginines relative to lysines in mediating DNA–core interactions reflects not only their comparable abundance but also the strategic positioning of these arginines at the DNA–histone interface, where they engage the DNA minor groove and form so-called anchoring contacts^1^. In contrast, the greater involvement of lysines over arginines in mediating tail–DNA interactions likely stems directly from their much higher abundance in the histone tails. Experiments have shown that histone-tail lysine acetylation destabilises nucleosome wrapping^27,50,51^. Consistently, our simulations show that both residue types are essential for maintaining a fully wrapped nucleosome: mutating all arginines to alanines or glutamines—while retaining lysines—leads to spontaneous unwrapping at zero force (Fig. S3A, B). A similar outcome is observed when all lysines are mutated, and arginines are left intact (Fig. S3C, D).

Among the four histones, the relative electrostatic contributions to nucleosome stability follow the trend H3*>*H2B*>*H2A*>*H4 for the core regions, and H3*>*H2B*>>*H4∼H2AN*>*H2AC for the histone tails (Fig. 2B). The distribution of charged residues within the core histones is: H3 (14 arginines, 5 lysines), H2A (9 arginines, 5 lysines), H2B (8 arginines, 12 lysines), and H4 (10 arginines, 6 lysines). In the flexible histone tails, the corresponding distributions are: H3 (4 arginines, 8 lysines), H2AN (3 arginines, 4 lysines), H2AC (0 arginines, 4 lysines), H2B (0 arginines, 8 lysines), and H4 (4 arginines, 5 lysines). Therefore, the relative electrostatic contributions of different histone regions to nucleosome stability reflect a combination of factors, including the number and positions of arginine residues at the histone core–DNA interface, the number and spatial distribution of charged residues within the histone tails (Fig. S4A), the tail lengths—which set their effective capture radii—and the locations at which the tails emerge from the nucleosome core (Fig. 2C).

Structural studies, atomistic simulations, and single-molecule experiments quantifying tail residence times and DNA–contact preferences report consistent trends^1,37,49,52–54^. Notably, our results agree with atomistic simulations of nucleosomes showing that the H3 tail has the longest residence time and strongest DNA binding among all tails, followed by H2B, H4, H2AN, and finally H2AC^54^. Further support for this hierarchy comes from additional atomistic simulations of nucleosomes, indicating that the H3 and H2B tails form the most stable contacts with DNA, whereas the H2AC tail is more dynamic and contributes less to DNA binding^54^. Our observations are also consistent with structural and biochemical experiments demonstrating that the H3 and H2B tails play roles in maintaining nucleosome stability^48,49,55^.

Crystallographic analyses show that the H2A C-tail interacts with nucleosomal DNA primarily near the dyad^56,57^. Although these contacts are weaker than those of H3 or H2B, biochemical experiments and simulations^52,58^ demonstrate that truncation of the H2A C-tail, or replacement with the shorter H2A.Z C-tail, increases nucleosome breathing^59^. This indicates that even a relatively modest change in tail composition at the dyad can influence nucleosome stability. This matches our simulations showing that the H2A C-tail forms the fewest contacts with DNA, yet still provides a small but measurable contribution to stabilising the wrapped state (Fig. 2C, S4D). Taken together, these findings support a hierarchy in which H3 and H2B dominate nucleosome stabilisation, while the H2A C-tail makes a weaker but locally important contribution.

When one of the outer-turn DNA arms detaches during stage 2, we observe the loss of several electrostatic contacts between DNA and charged residues within the H3 and H2A histone cores (Stage 2; Fig. 2B, S4B, D). As H3 and H2A are positioned adjacent to the nucleosomal DNA entry-exit region, they are the first core histones to lose contact with DNA upon partial unwrapping (Fig.2C, Supplementary Video 1). Notably, however, interactions between charged residues in the histone tails and DNA remain largely intact during this stage. This suggests that tail–DNA contacts function as stabilising anchors that withstand higher forces than the corresponding core–DNA contacts, thereby helping preserve nucleosome integrity under moderate tension. Recent simulations^48,49,53^ and single-molecule experiments^39,40^ have similarly proposed that, by maintaining these local electrostatic contacts, the tails not only stabilise partially unwrapped states but can also facilitate DNA rewrapping once the tension is released. In this view, the tails act as flexible guides that promote recovery of the fully wrapped nucleosome conformation.

As the applied force increases during full outer-turn unwrapping (Stage 3), we observe a pronounced loss of electrostatic contacts between core residues and DNA—again most prominently for H3 and H2A, consistent with their proximity to the entry–exit DNA region (Fig.2B, S4B, D). Concurrently, tail–DNA interactions, particularly those involving the H3 tail and, to a lesser extent, the H4 tail, also diminish substantially (Fig.2B, S4C). This progressive loss of stabilising associative histone–DNA contacts reflects the increasing mechanical stress placed on the nucleosome, further reinforcing the idea that core and tail residues play coordinated roles in resisting unwrapping as the force increases.

During unwrapping of the inner turn (Stage 4), the remaining electrostatic interactions between histone residues and DNA—both from the core and the tails—are progressively lost. Notably, contacts involving H2B residues persist the longest, likely due to the positioning of H2B toward the nucleosome back, indicating that H2B residues may help delay complete DNA release in the final stages of unwrapping (Fig. 2 B, C, S4E, Supplementary Video 1). In our simulations, once full unwrapping is achieved, we observe an intermediate state in which the intact histone core remains associated with the extended DNA (Fig. 2C). However, under experimental conditions, this intermediate would likely be bypassed altogether, with complete nucleosome disassembly occurring instead^4,60^.

In summary, when comparing the respective roles of the different histone charged residues to nucleosome stability, two important trends emerge: (1) histone core–DNA interactions, particularly of H3 and H2A, break down first (i.e., at lower forces) and are primarily mediated by arginines, and (2) histone tail–DNA interactions resist higher forces and are mostly contributed by lysines, predominantly of H3 and H2B. The histone tail–DNA contacts are likely more resilient due to the intrinsic disorder and flexibility of the tails, which can adapt their conformations to maintain contact with the DNA under the diverse configurations imposed by mechanical stress.

### Histone modifications can significantly alter nucleosome stability against applied force

In this section, we examine how changes in histone composition influence the free energy landscape of nucleosome unwrapping by performing additional umbrella sampling simulations and computing PMFs for nucleosomes with histone variants and different patterns of lysine acetylation, one of the most prevalent chromatin marks. To quantitatively compare the impact of the modifications on the unwrapping response of the nucleosome, we compute the equilibrium free-energy difference between the fully wrapped and fully unwrapped states under tension. This quantity is obtained as the difference between the values of the PMF along the extension coordinate, ΔΔ*G* = Δ*G*(*x*_unwrapped_) − Δ*G*(*x*_wrapped_), where Δ*G*(*x*) is the PMF value and *x*_wrapped_ and *x*_unwrapped_ are the reaction coordinate values at which the nucleosome is structurally classified as fully wrapped and fully unwrapped, respectively. The value of ΔΔ*G* corresponds to the reversible work required to convert a wrapped nucleosome into its fully unwrapped state under force. Therefore, smaller values indicate a nucleosome that is more plastic (i.e., more mechanically compliant), whereas larger values correspond to greater mechanical resistance to unwrapping and lower nucleosome plasticity.

In an experimental pulling assay, the measured mechanical work would, in general, correspond to irreversible work and exceed this value due to energy dissipation. All numerical values for ΔΔ*G*^mod^ for the modified systems and their absolute differences with respect to the canonical nucleosome are provided in Supplementary Table 2.

Because acetylation patterns vary widely across nucleosomes and cell types^61^, we performed a systematic assessment of how the location and extent of acetylation modulate nucleosome stability. We therefore consider five classes of modifications: (i) hyperacetylation of lysines in the core and tails; (ii) hyperacetylation of core lysines; (iii) hyperacetylation of tail lysines; (iv) acetylation of all lysines in the H3 tail and H2A C-terminal tail; and (v) experimentally observed endogenous acetylation patterns (H3K9ac, H3K14ac, H3K18ac, H3K23ac, H4K5ac, H4K8ac, H4K12ac, H4K16ac and H4K20ac)^62^ (Fig. 3A, Supplementary Table 2).

**Figure 3.**
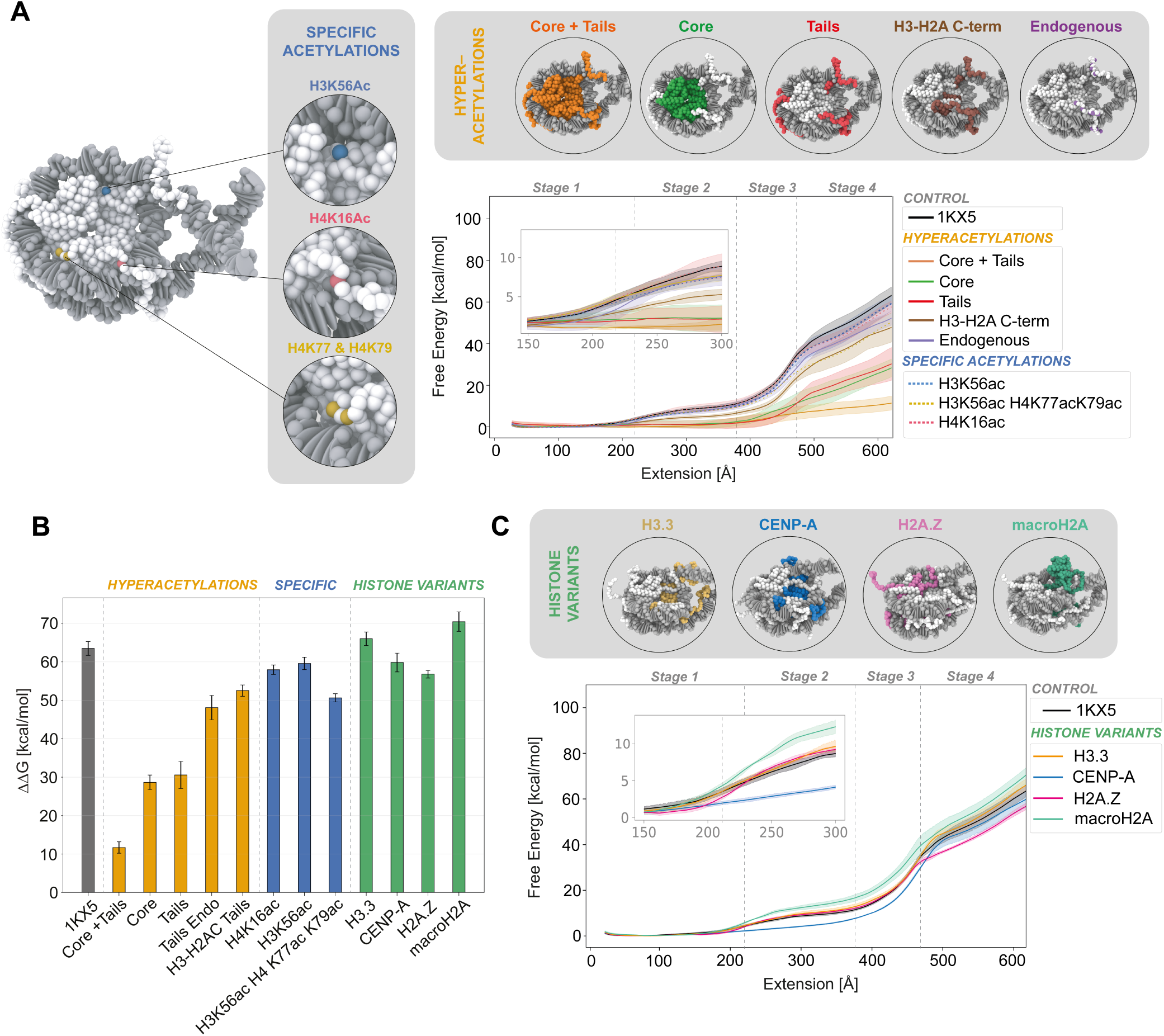
Histone modifications differentially facilitate force-induced nucleosome unwrapping. **A.** Structural representation of the selected acetylation patterns (left and top) and corresponding free-energy profiles (right). Global acetylation schemes (all Lys, core Lys, tail Lys, H3–H2A C-terminal Lys, endogenous Lys) and specific acetylations (H3K56ac, H3K56ac/H4K77ac/H4K79ac, H4K16ac) are shown. **B.** ΔΔ*G* of nucleosome unwrapping evaluated at an extension of 620 Å relative to the fully wrapped nucleosome. Bars show mean ± s.e.m. **C.** Models of nucleosomes incorporating histone variants, with CENP-A highlighted in blue, H3.3 in yellow, H2A.Z in pink, and macroH2A in green (top). Corresponding force–extension curves for the indicated histone variants are shown below, using the same color scheme.

Extensive lysine acetylation produces a pronounced increase in nucleosome plasticity, significantly facilitating force-induced unwrapping (Fig.3B). Hyperacetylation of lysines in both the histone core and tails leads to a dramatic reduction in the free-energy difference between the wrapped and unwrapped states (−51.81 ± 2.35 kcal/mol) relative to the canonical nucleosome (Fig.3B and Supplementary Table 2).

Decomposition of the electrostatic interactions provides a mechanistic basis for these trends. Analysis of the Debye–Hückel electrostatic energy, resolved by residue type, histone type, and unwrapping stage (Fig. S5), reveals that hyperacetylation of tail lysines produces a near-complete loss of the dominant tail-mediated electrostatic interactions that, in the canonical nucleosome (Fig. 2), help maintain histone–DNA contacts in partially unwrapped states. In contrast, acetylation of core lysines suppresses direct lysine–DNA attraction in the core and additionally induces a substantial indirect reduction in core arginine–DNA electrostatics, reflecting the reduced ability of DNA to remain associated with the histone core once lysine-mediated anchoring is weakened (Fig. S5). When both tail and core lysines are acetylated simultaneously, electrostatic stabilisation is strongly diminished across the entire nucleosome, leaving residual arginine-mediated interactions that are insufficient to sustain DNA wrapping under force (Fig. S5).

Restricting hyperacetylation to either the histone core or the histone tails likewise results in substantial enhancement of nucleosome plasticity, albeit with distinct signatures (Fig. S5). Core acetylation decreases ΔΔ*G* by 34.85 ± 2.63 kcal/mol relative to the canonical case, whereas tail acetylation yields a reduction of 32.93 ± 3.96 kcal/mol. As the number of acetylated residues is further reduced, nucleosome plasticity remains enhanced relative to the canonical state, although the magnitude of the effect progressively decreases. For instance, acetylation restricted to the H3 tail and the H2A C-terminal tail, both localised near the dyad, lowers ΔΔ*G* by 15.39 ± 3.64 kcal/mol (Fig. 3B).

Physiologically relevant acetylation patterns produce subtler but mechanistically informative effects, demonstrating that the enhancement of nucleosome plasticity is not determined solely by the total number of acetylated lysines (Fig. 3B). For instance, we investigate the acetylation of H4K16, which is the classical mark of accessible chromatin^27^, and the acetylation of H3K56ac, H4K77ac, and H4K79ac, which are associated with increased DNA accessibility and nucleosome unwrapping^63^. Single-site acetylation of H4K16 or H3K56 alone produces comparatively small effects, decreasing ΔΔ*G* by 5.53 ± 2.18 and 3.90 ± 2.42 kcal/mol, respectively, indicating only a modest increase in nucleosome plasticity. Combined acetylation of H3K56, H4K77, and H4K79 reduces ΔΔ*G* by 12.86 ± 2.10 kcal/mol (Fig. 3B). Strikingly, the effect of these three combined acetylations is comparable to that observed when all nine known endogenous acetylation sites (H3K9ac, H3K14ac, H3K18ac, H3K23ac, H4K5ac, H4K8ac, H4K12ac, H4K16ac and H4K20ac) are modified simultaneously (Fig. 3B). This non-additive behaviour demonstrates that nucleosome unwrapping is sensitive not simply to the total degree of acetylation, but to the specific identity, spatial arrangement, and interactions between acetylation marks.

Overall, the dominant effect of acetylation is a substantial reduction in the forces required to unwrap the nucleosome, as expected from weakening of DNA–histone contacts (Fig. S5). For acetylation patterns involving only a small number of sites, the specific identity and positioning of the modified residues dominate their impact on increasing nucleosome plasticity, leading to non-additive modulation of nucleosome plasticity. In contrast, under extensive hyperacetylation, where lysine-mediated electrostatic interactions are globally suppressed, the enhancement of nucleosome plasticity becomes largely cumulative. In both regimes, neutralisation of lysines removes their direct electrostatic attraction to DNA and compromises the ability of adjacent arginine residues to maintain persistent DNA contacts, thereby weakening the collective histone–DNA contact network that underpins mechanically protected unwrapping intermediates.

We next focus on histone variants, which provide an alternative regulatory mechanism by altering the chemical composition and structure of the histone core. To test the impact of replacing canonical histones with variants found *in vivo*, we compare the effects of H3.3, CENP-A, H2A.Z, and macroH2A (Fig. 3C, Supplementary Table 2). Overall, these histone variants produce mild to moderate changes in nucleosomal stability relative to the canonical nucleosome, consistent with their roles in fine-tuning chromatin structure rather than globally disrupting nucleosome integrity.

The H3.3 variant differs from canonical H3 by only four amino acid substitutions (A31S, S87A, I89V, and A96S), all of which occur in non-charged residues. Accordingly, this minimal chemical perturbation of the histone core results in no remarkable change in nucleosome plasticity with ΔΔ*G* differing by 2.53 ± 2.53 kcal/mol relative to the canonical octamer (Fig. 3C).

CENP-A shares approximately 60% sequence identity with canonical H3 but, importantly, possesses a substantially shorter N-terminal tail^64^. Incorporation of CENP-A decreases ΔΔ*G* by 3.66 ± 3.02 kcal/mol relative to the canonical nucleosome (Fig.3C), corresponding to a modest increase in nucleosome plasticity under tension. Mechanistically, this enhancement arises primarily from a pronounced reduction of the first mechanical barrier, which decreases by approximately 58%, while the second barrier remains largely unchanged (Fig.3C; Supplementary Table 4). Thus, CENP-A selectively increases outer-turn plasticity while largely preserving the resistance associated with inner-turn DNA contacts.

Together, these features indicate that CENP-A nucleosomes are mechanically more plastic than their canonical counterparts. This interpretation is consistent with previous reports of increased DNA-end flexibility in CENP-A nucleosomes^65^, and with the observation that CENP-A nucleosomes at centromeres—assembled on repetitive DNA sequences lacking strong positioning motifs^66^—form more dynamic, fuzzy nucleosomes capable of repositioning readily^67^.

The H2A variant H2A.Z shares approximately 60% sequence identity with canonical H2A and possesses a shorter C-terminal tail. Previous simulations have suggested that incorporation of H2A.Z weakens DNA–histone interactions at the entry–exit regions, thereby enhancing nucleosome breathing^59^. Consistent with this picture, we find that H2A.Z decreases ΔΔ*G* by 6.70 ± 2.08 kcal/mol relative to the canonical nucleosome (Fig. 3C), corresponding to a moderate increase in nucleosome plasticity under tension. Mechanistically, this enhancement arises from differential modulation of the two mechanical unwrapping barriers: H2A.Z increases the first barrier by approximately 36%, but reduces the second by about 20% (Fig.3C; Supplementary Table 4). Because the second barrier constitutes the dominant mechanical bottleneck and accounts for the largest contribution to the free-energy difference along the unwrapping pathway, its reduction outweighs the increased resistance of the outer turn. This redistribution of barrier heights is reflected in the PMF (Fig. 3C) and results in an overall moderate increase in nucleosome plasticity.

Lastly, macroH2A—an H2A variant containing an additional globular domain—produces the opposite effect, modestly reducing nucleosome plasticity under tension. Incorporation of macroH2A increases ΔΔ*G* by 6.98 ± 3.10 kcal/mol relative to the canonical nucleosome (Fig.3C), indicating a larger free-energy difference between wrapped and unwrapped states. This effect is explained by a pronounced increase in the first mechanical barrier for unwrapping, which rises by approximately 73% (Fig.3C; Supplementary Table 4). These findings align with the established role of macroH2A in promoting compact and mechanically resilient chromatin domains, such as those associated with X-chromosome inactivation^68^.

Taken together, these results demonstrate that histone variants modulate nucleosome mechanics through localised and chemically specific perturbations of the histone–DNA interaction network. In contrast to lysine acetylation, which invariably weakens histone–DNA electrostatic interactions and increases nucleosome plasticity, hisitone variants exert a bivalent regulatory effect: depending on the nature and location of the substitution, they can either increase nucleosome plasticity, as observed for CENP-A and H2A.Z, or decrease nucleosome plasticity by reinforcing DNA–histone anchoring, as in the case of macroH2A. This highlights the essential role of chemical heterogeneity in regulating how nucleosomes respond to mechanical stress, for instance, during remodelling.

### DNA sequence sets the balance between nucleosome stability and mechanical plasticity

We next examine how changes in the DNA sequence modulate the unwrapping response of nucleosomes under force (Fig.4A–E). To this end, we compare the canonical nucleosome with the 1KX5 sequence with other strong nucleosome-positioning sequences, a set of synthetic DNA sequences, and several biologically relevant genomic sequences (Supplementary Table 3).

**Figure 4.**
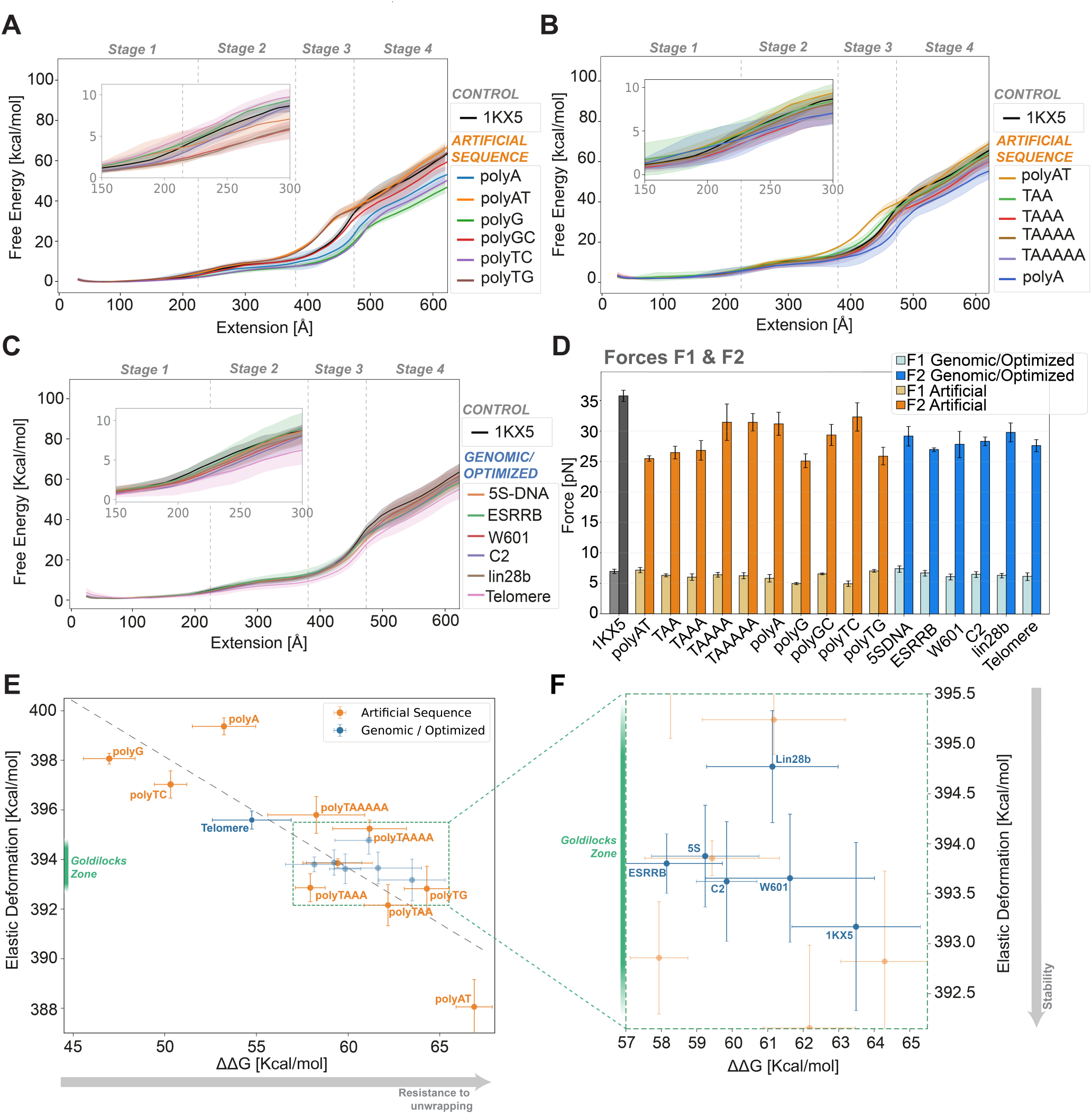
DNA sequence sets a trade-off between nucleosome thermodynamic stability and nucleosome plasticity. **A.** Free-energy profiles (PMFs) for nucleosomes assembled on artificial repetitive DNA sequences (polyA, polyAT, polyG, polyGC, polyTC, polyTG). Shaded regions indicate standard errors. **B.** PMFs for shorter artificial repeats (TAA, TAAA, TAAAA, TAAAAA). **C.** PMFs for nucleosomes assembled on native sequences (5S-DNA, ESRRB, W601, Lin28b, C2, telomeric DNA). **D.** Forces associated with the first (F1) and second (F2) unwrapping barriers for artificial (orange) and genomic/optimized (blue) sequences. Bars show mean ± s.e.m. **E-F.** ΔΔ*G*_620_ of nucleosome unwrapping versus elastic deformation energy of wrapped DNA. Lower ΔΔ*G*_620_ indicates increased mechanical plasticity and easier unwrapping under force, whereas higher deformation energy is associated with reduced nucleosome stability. Artificial (orange) and genomic/optimized (blue) sequences follow distinct trends, with artificial sequences showing a strong correlation (*R*^2^ = 0.7). Boxed region indicates the sequences within a ‘Goldilocks zone’ of intermediate DNA elastic deformation energy.

To mechanistically rationalize the impact of DNA sequence-dependent mechanical properties on the response of nucleosomes to unwrap, we compute the elastic deformation energy of the DNA (see Methods and Figs. 4E–F and S6). This deformation energy provides a sequence-dependent measure of the intrinsic energy required to bend and twist the DNA for nucleosomal wrapping and thus offers a physical interpretation of the observed variability in mechanical stability. The change in free energy between fully wrapped and fully unwrapped states (ΔΔ*G*) exhibits a strong inverse correlation with the intrinsic mechanical deformation energy of each DNA sequence (Fig.4E–F), reflecting the monotonic contribution of DNA elasticity to the mechanical cost of unwrapping. We note that this quantity characterises the mechanical resistance of an already assembled nucleosome to force-induced unwrapping rather than the propensity of nucleosome formation. Specifically, ΔΔ*G* is the equilibrium free-energy difference between the wrapped and unwrapped states under the applied force. Thus, ΔΔ*G* should not be interpreted as a measure of the propensity of a DNA sequence to form or position nucleosomes.

We tested ten different synthetic repetitive DNA sequences (Fig.4A–C), all of which modulate the mechanical response of nucleosomes by either increasing or decreasing the free-energy difference between wrapped and unwrapped states (ΔΔ*G*) relative to the 1KX5 nucleosome (Fig.4E–F). Among these DNA sequences, polyG, polyA and polyTC exhibit the highest DNA deformation energies (i.e., they are stiffer), and consequently display the lowest ΔΔ*G* values, corresponding to the greatest mechanical plasticity and the lowest resistance to force-induced unwrapping (Fig. 4E–F). For polyG, this increase in nucleosome plasticity is also reflected in a substantial reduction of both unwrapping force barriers (Δ%*F*_1_ = −28.8 ± 4.5%, Δ%*F*_2_ = −29.8 ± 3.7%), whereas for polyA and polyTC the reductions are more modest (Fig. 4D). At the opposite extreme (i.e., the most flexible sequences), polyAT exhibits the lowest deformation energy and, as a result, the largest ΔΔ*G*, indicating the greatest resistance to force-induced unwrapping and therefore the lowest mechanical plasticity. PolyTG follows the same qualitative trend, albeit less strongly, exhibiting the second-lowest deformation energy and the second-largest ΔΔ*G*. Strikingly, despite their increased resistance to complete unwrapping, both sequences exhibit a substantial reduction in the second force barrier (Δ%*F*_2_ = −28.8 ± 2.2% and −27.6 ± 4.4%, respectively), while leaving the first barrier essentially unchanged relative to the canonical sequence.

This behaviour initially appears counter-intuitive: the most flexible sequences (polyAT, polyTG) yield the nucleosomes that require the most total work to fully unwrap (largest ΔΔ*G*), yet display the lowest unwrapping peak forces. To understand how increased flexibility raises the integrated work despite lowering the peak force, we examined how DNA mechanics modulates not only the height of the force barriers but also the emergence of the topologically protected partially unwrapped intermediates along the extension coordinate. We focus on the second intermediate, as it is associated with the highest force barrier, and hence the true bottleneck of the unwrapping process. To do so, we quantified the shift in the extension at which the second force maximum occurs, Δ*X*_F2_, relative to the canonical nucleosome (Fig.S6D). Negative values of Δ*X*_F2_ indicate earlier emergence of this intermediate and appear consistently for DNA sequences with increased flexibility, whereas positive values occur for stiffer sequences, for which DNA configurations are more mechanically constrained (Fig. 4E–F).

For highly flexible sequences such as polyAT and polyTG, the second topologically protected intermediate appears at substantially smaller extensions (Δ*X*_F2_ *<* 0; Fig.S6D). Importantly, the increased flexibility of these sequences not only facilitates earlier access to the intermediate state but also over-stabilises it over a broad range of extensions, allowing the nucleosome to sustain a partially unwrapped, topologically protected configuration for longer, despite a reduced force barrier. As a consequence, although the peak force associated with the second transition is lower, the force remains elevated over a wider range of extensions, leading to a remarkable increase in the total pulling work required for unwrapping. In contrast, stiff sequences such as polyG, polyTC and polyA shift the second barrier to larger extensions (Δ*X*_F2_ *>* 0; Fig.S6D), reflecting the higher resistance of the rigid DNA sequences to deformation. In this case, DNA rigidity not only delays access to the partially unwrapped intermediate but also reduces its persistence—and that of the associated elevated force values—to a narrower range of extensions, resulting in a much lower overall free-energy difference between wrapped and unwrapped states than would be inferred from the modest reduction in barrier heights alone.

Overall, the correlations among Δ*X*_F2_, elastic deformation energy (polyA, polyG, polyTC ≫ polyGC, 1KX5 ≫ polyTG, polyAT; Fig. S6), and ΔΔ*G* demonstrates that DNA mechanics controls ΔΔ*G* through both the heights of the force barriers and the stability and persistence of the topologically protected intermediates along the extension coordinate. Consequently, ΔΔ*G* is larger for flexible sequences because those sequences sustain the topologically protected intermediates over a broader range of extensions, whereas stiffer sequences progress more readily to the fully unwrapped state despite exhibiting higher force barriers.

To probe the role of local sequence periodicity, we simulated a series of A-tract variants with stepwise increases in TA repeat length (polyAT, TAA, TAAA, TAAAA, TAAAAA) (Fig.4B–F and S6). These sequences reveal a clear trend linking DNA flexibility to nucleosome stability and plasticity. The shortest-period sequence, polyAT, which is the most flexible, exhibits the largest free-energy difference between wrapped and unwrapped states, and hence the greatest resistance to complete force-induced unwrapping, while simultaneously exhibiting a markedly earlier onset of the second unwrapping transition (Δ*X*_F2_ *<* 0). Consistent with the analysis above, this early emergence is accompanied by an increased persistence of the partially unwrapped intermediate over a broad range of extensions, thereby increasing the cumulative mechanical work required for complete unwrapping of polyAT nucleosomes. As the A-tract length increases and DNA flexibility decreases, the free-energy difference between wrapped and unwrapped states progressively decreases (indicating increased plasticity), and the position of the second force barrier shifts towards the 1KX5 value (Δ*X*_F2_ → 0). For the fully A-tract sequence, polyA, the second barrier occurs at larger extensions (positive Δ*X*_F2_; Fig.S6D) and the free-energy difference is comparatively low, consistent with reduced resistance to force-induced unwrapping.

The periodic arrangement of A/T- and G/C-rich motifs relative to the ∼10-bp DNA helical repeat is well known to modulate DNA curvature, bendability, and sequence-dependent nucleosome positioning^69^. This effect is reflected in the TAAAA repeat, which exhibits a larger ΔΔ*G* than both the TAAA and TAAAAA sequences (Fig.4E-F). Because the 5-bp TAAAA repeat corresponds approximately to half of the ∼10-bp DNA helical repeat, successive A-tracts realign on the same helical face once per full turn (i.e. every two repeats), so their intrinsic bending preferences accumulate coherently and can enhance DNA curvature. The 4-bp and 6-bp periods phase less cleanly with the helical repeat, which is consistent with the smaller ΔΔ*G* of polyTAAA and polyTAAAAA. By contrast, PolyAT and PolyTAA, with the shortest A-runs and hence the greatest flexibility of the series, retain a large ΔΔ*G* despite incoherent phasing. In addition, the increase in ΔΔ*G* for the TAAAA repeat is modest compared with the overall variation in ΔΔ*G* across the A-tract series. These results indicate that DNA flexibility establishes the dominant trend in ΔΔ*G*, while sequence phasing provides a secondary modulation. Previous all-atom simulations have similarly shown that sequence periodicity in phase with the DNA helix amplifies intrinsic DNA curvature, whereas out-of-phase variants reduce this effect^70^.

We next compare the behaviour of the 1KX5 sequence with three additional strong nucleosome-positioning sequences: the Widom 601 sequence (W601)^69^, its derivative C2^71^, and the naturally occurring 5S DNA sequence derived from the highly expressed 5S ribosomal RNA gene locus^72,73^. Unlike the synthetic repeats, these sequences share the canonical ∼10-bp periodic arrangement of A/T- and G/C-rich motifs in anti-phase, which positions the flexible A/T-rich motifs where the DNA minor groove faces the histone octamer, while the stiffer G/C-rich motifs occupy the opposing orientation, thereby favouring the preferred rotational orientation of DNA around the histone octamer. All four sequences exhibit comparable free-energy differences between the wrapped and unwrapped states (Fig. 4E–F), together with similar force barriers and a similar onset of the partially unwrapped intermediates (Fig. 4D and Fig. S6D), consistent with their ability to form stable, well-positioned nucleosomes and with previous experimental observations^31,39–42^.

The W601 sequence was selected by SELEX for high-affinity, precise nucleosome positioning^69^. Nonetheless, W601 and the other strong positioners cluster within the intermediate range of elastic deformation energies and do not reach the maximal resistance to force-induced unwrapping (highest ΔΔ*G*) displayed by the artificial polyAT sequence (Fig.4E–F). We recall that ΔΔ*G* reports the mechanical resistance of an already assembled nucleosome to force-induced unwrapping, rather than the thermodynamic affinity for nucleosome formation. The highest ΔΔ*G* value arises for the highly deformable polyAT because it over-stabilises the partially unwrapped intermediates. By contrast, efficient nucleosome formation and positioning require a balance between DNA deformability and sequence patterning. Specifically, the DNA must be deformable enough to wrap at moderate energetic cost around the histone core, yet must also retain the proper periodicity of A/T- and G/C-rich motifs. Achieving this periodic sequence pattern generally compromises maximal deformability because it interleaves highly flexible A/T-rich motifs with mechanically stiffer G/C-rich motifs. As a result, strong-positioning sequences naturally occupy an intermediate region of deformation energy rather than the extreme represented by polyAT. By contrast, polyAT attains the largest ΔΔ*G* in our set because its exceptionally low deformation energy over-stabilises partially unwrapped intermediates across a broad range of extensions, increasing the total mechanical work required for complete unwrapping, while lacking the sequence pattern needed for strong nucleosome positioning.

We note one limitation of our approach relevant to these strongly patterned sequences. Because our model represents DNA using the sequence-dependent Rigid Base Pair model^74^, and therefore does not resolve a physical minor groove, it omits the A/T minor-groove narrowing that, in atomistic structures, enhances arginine insertion into the minor groove and strengthens histone–DNA contacts. Thus, the ΔΔ*G* of these sequences may be modestly underestimated. We expect this effect to be small, however, because our model already captures the dominant histone–DNA electrostatic anchoring—including the arginine–DNA interactions discussed in the previous section. Therefore, the explicit contribution of minor-groove narrowing is likely to provide only an incremental correction and would not alter the conclusion that these sequences remain within the intermediate deformation—energy range and below the maximal-resistance polyAT extreme.

Beyond these strong positioning sequences, we examined DNA drawn from genomic contexts associated with distinct chromatin environments, including nucleosomes positioned at the promoters of the pluripotency genes Lin28b and ESRRB^75,76^ and a repetitive telomeric sequence known to form specialised chromatin at chromosome ends^77,78^ (Telomere in the figures). Notably, all these nucleosome-forming biological DNA sequences show free-energy difference between wrapped and unwrapped states and values of the height and location of the unwrapping force barries comparable to one another and to those of the strong-positioning sequences (Fig. 4C–F). Their similar wrapping energetics is consistent with experimental evidence showing that all of these sequences form stable nucleosomes either *in vitro*^37,71^ or *in vivo*^75,76^. We further analysed DNA sequences derived from *in vivo* nucleosome-depleted regions (NDRs), nucleosome-positioning sequences (NPSs), and random non-periodic sequences (Fig. S6). As expected, the NDR sequence exhibits both high elastic deformation energy and a comparatively small

ΔΔ*G*, consistent with its poor ability to form stable nucleosomes *in vivo*. By contrast, the NPS exhibits intermediate values of both elastic deformation energy and ΔΔ*G*, comparable to those of the strong-positioning and genomic sequences. We note that there is no experimental evidence that the random non-periodic sequences we have analyzed form stable nucleosomes *in vitro*. Nevertheless, when simulated in the nucleosomal state, these random sequences display the same inverse relationship between elastic deformation energy and ΔΔ*G*, further supporting the conclusion that DNA deformability is a major determinant of the mechanical resistance of assembled nucleosomes to force-induced unwrapping.

A fascinating result stemming from our work is that both genomic and strong nucleosome-positioning sequences cluster within a comparatively narrow range of intermediate deformation energies and nucleosome plasticity, as inferred from the free-energy difference between wrapped and unwrapped states (boxed region in Fig.4E–F). Importantly, this regime does not correspond to a maximum of resistance to force-induced unwrapping, but instead defines a mechanically permissive range in which nucleosomes are thermodynamically stable while retaining sufficient plasticity to undergo controlled deformation under applied force. DNA sequences that are too mechanically rigid incur a large elastic deformation penalty upon wrapping, destabilising the fully wrapped state. Conversely, excessively flexible DNA increases nucleosome stability but reduces the mechanical plasticity of the nucleosome by over-stabilising wrapped and partially unwrapped configurations, including topologically protected intermediates over a larger range of extensions (see Fig. S8). We therefore propose that this intermediate regime constitutes a ‘Goldilocks zone’ of DNA flexibility, in which nucleosome stability and nucleosome plasticity are optimally balanced.

### Synergistic modulation of nucleosome unwrapping by DNA sequence and histone composition

We next investigated how simultaneous changes in histone composition and DNA sequence can produce cumulative or non-additive effects on nucleosome mechanics (Supplementary Table 4). Such effects may not be apparent when each component is examined in isolation.

As a case study, we analysed the interplay between the centromeric histone variant CENP-A and the alpha satellite DNA sequence. Both are enriched at centromeres and have been proposed to act synergistically to specify centromeric chromatin^79,80^ (Fig. 5A). We compared the PMFs of four nucleosome constructs: (i) the canonical 1KX5 nucleosome, (ii) 1KX5 histones with the alpha-satellite DNA, (iii) 1KX5 DNA with the H3 histone replaced by CENP-A, and (iv) the fully centromeric-like construct combining both CENP-A and alpha-satellite DNA.

**Figure 5.**
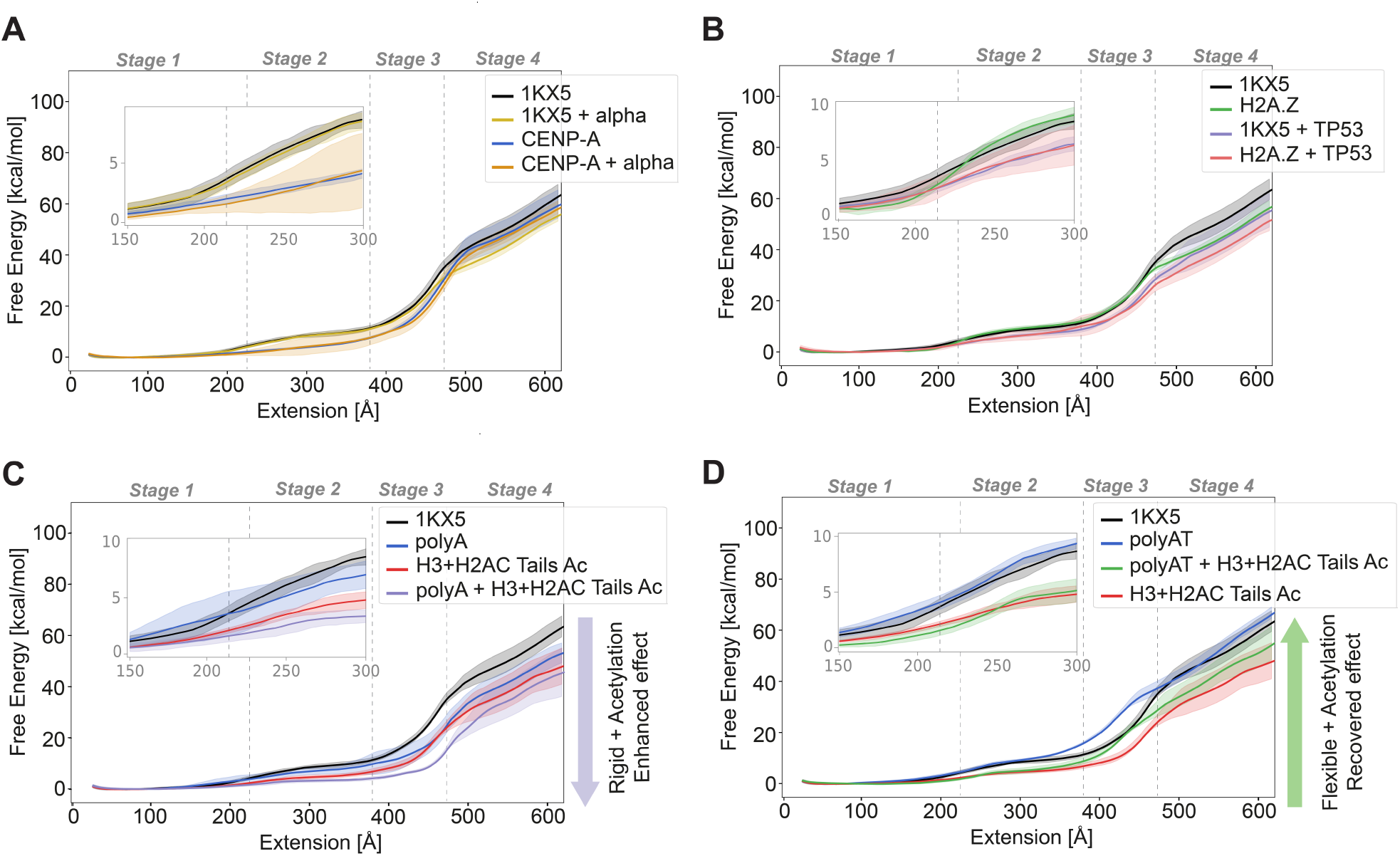
Combined changes in DNA sequence and histone composition act cooperatively to regulate nucleosome responses to force. **A.** Pulling curves for the 1KX5 and CENP-A histone nucleosomes, with the 1KX5 sequence and the genomic alpha satellite sequence. **B.** Pulling curves for the 1KX5 and H2A.Z histone nucleosomes, with the 1KX5 sequence and the TP53 sequence. **C.** Pulling curves of the canonical 1KX5 nucleosome and incorporating a polyA sequence, with and without the H3/H2A c-term tails acetylation pattern. **D.** Pulling curves of the canonical 1KX5 nucleosome and incorporating a polyAT sequence, with and without the H3/H2A C-term tails acetylation pattern.

Incorporation of CENP-A, whether paired with optimized DNA or with genomic alpha-satellite DNA, attenuates the outer-turn unwrapping barrier. This behaviour is consistent with CENP-A weakening histone–DNA interactions near the entry and exit sites. In contrast, pairing genomic alpha-satellite DNA with canonical histones specifically lowers the free-energy barrier associated with inner-turn unwrapping (Fig. 5A), which is reflected in a significantly reduced *F*_2_ peak (Fig. S7A).

Strikingly, when CENP-A and genomic alpha-satellite DNA are combined, their effects are not additive. Although CENP-A continues to weaken the early unwrapping barrier, the reduction in the second barrier observed for alpha-satellite DNA plus canonical histones is no longer present (Fig. 5A). Instead, the fully centromeric construct displays Δ*X*_F2_ values that are almost indistinguishable from those of the canonical nucleosome (Fig. S7B). This indicates that CENP-A counterbalances the destabilising influence of the genomic alpha-satellite sequence on inner-turn unwrapping, restoring the mechanical stability of the inner turn to a canonical-like level.

For the case of H2A.Z, we used the TP53 promoter DNA sequence, as this histone type is typically enriched at promoters^81^. The TP53 promoter is characterised by dynamic chromatin architecture and high nucleosome turnover^82^, and H2A.Z enrichment at this site has been linked to enhanced transcriptional activation through modulation of nucleosome stability and accessibility^83–85^. Consistent with this view, incorporating TP53 DNA into the canonical nucleosome lowers its unwrapping free energy and reduces both the first and second force barriers, indicating a destabilising effect (Fig. 5B). Replacing H2A with H2A.Z, while keeping the 1KX5 DNA, increases the value of *F*_1_ but lowers the *F*_2_ value (Fig. S7A). When H2A.Z and TP53 DNA are combined, their effects are cumulative: both unwrapping barriers are reduced, and *F*_2_ decreases even further than in the TP53–canonical histones nucleosome. The H2A.Z–TP53 nucleosome is therefore substantially more labile than either single-component mutant, indicating that, unlike the compensatory behaviour observed for CENP-A and genomic alpha-satellite DNA, H2A.Z and TP53 DNA act cooperatively to destabilise the nucleosome.

Finally, to dissect how the mechanical response of nucleosomes depends on the interplay between histone modifications and DNA rigidity, we tested whether acetylation of the H3 tail and H2A C-terminal tail (which reduces the strength of electrostatic histone–DNA interactions) could produce cumulative effects when combined with DNA sequences with increased or decreased flexibility.

To this end, we first combined histones with the H3 and H2A C-terminal tail acetylated with a rigid DNA sequence (polyA), which by itself also reduces stability due to its resistance to bending. This combination resulted in a pronounced decrease in mechanical stability (Fig. 5C), fully flattening the outer-turn barrier and significantly lowering the inner turn barrier, confirming that histone and DNA characteristics can synergize to amplify the unwrapping propensity.

Conversely, when the same acetylated histones were paired with a highly flexible sequence (polyAT), we observed a partial recovery of nucleosome mechanical resistance during inner-turn unwrapping (Fig. 5D), while the outer-turn barrier remained suppressed. This suggests that DNA flexibility can compensate for destabilising histone modifications, restoring unwrapping resistance by reducing the cost of bending DNA to wrap around the histone.

Together, these results demonstrate that histone modifications and DNA mechanics function cooperatively, and that the net mechanical response of a nucleosome arises from the context-dependent interplay between chemical composition, DNA mechanics, and structural features of both histones and DNA.

## Discussion

In this work, we provide a comprehensive biophysical framework for understanding the thermodynamic and mechanical factors that govern nucleosome stability under force. Using equilibrium umbrella sampling simulations with a near-atomistic, chemically specific coarse-grained chromatin model, we systematically compare force-extension and free energy curves for more than 40 nucleosomal systems, varying in DNA sequence—both synthetic and genomic—histone variants, and acetylation patterns. Our simulations reveal that DNA sequence and histone chemistry regulate nucleosome unwrapping on markedly different scales: whereas sequence-dependent DNA mechanics tunes the unwrapping response within a relatively narrow regime, changes in histone composition and post-translational modifications produce substantially larger changes in the force barriers and the total reversible work of unwrapping (ΔΔ*G*).

We find that the characteristic force barriers observed during nucleosome unwrapping emerge from a complex interplay between histone–DNA and DNA–DNA electrostatic interactions, sequence-dependent DNA mechanics, and nucleosome geometry. In particular, we identify two prominent force peaks in the pulling curves, also observed experimentally^40,41,63,86,87^, which correspond to the disruption of distinct, topologically protected intermediate states (Supplementary Table 1). In these topologically protected states, the applied force is misaligned with the DNA path, and therefore cannot efficiently drive unwrapping. Breaking out of these protected states requires reorienting the nucleosome and aligning the DNA path to the pulling direction. Indeed, we find that histone–DNA contacts break easily once the force overcomes the protected configuration, suggesting that these maxima are best interpreted as thresholds for releasing protected configurations, rather than for fully unwrapping the outer and inner turns.

Consistently, we find that the enhanced thermodynamic stability of nucleosomes assembled on highly flexible DNA arises from the stabilisation of the topologically protected, partially unwrapped intermediates encountered during unwrapping. Highly flexible sequences such as polyAT and polyTG allow the sharply bent DNA conformations within these intermediates to remain stable over a broad range of extensions before the DNA path reorients into a configuration from which unwrapping can proceed. As a result, formation of these intermediates occurs at a lower peak force, even though such intermediates, and hence the elevated-force regime, persist over a longer extension range. Indeed, highly flexible sequences exhibit smaller values of the second force maximum (*F*_2_) while requiring greater total reversible work for complete unwrapping, because ΔΔ*G* reflects the area under the force–extension curve rather than the peak force alone. Conversely, more rigid sequences such as polyG, polyA and polyTC pay a larger energetic cost to bend around the histone core, destabilising both the wrapped state and the topologically protected intermediates. These intermediates therefore persist over a narrower extension range and at higher peak forces, giving rise to mechanically more plastic nucleosomes that nevertheless require less total work to unwrap completely.

Strikingly, we find that both genomic and strong nucleosome-positioning sequences—i.e., ESRRB, Lin28b, the TP53 regulatory region, 1KX5, 5S, W601, and C2—cluster within a narrow ‘Goldilocks’ regime of intermediate DNA flexibility. DNA that is too flexible incurs a low energetic cost of bending around the histone core, but limits the mechanical responsiveness of the nucleosome. Conversely, DNA that is too rigid incurs a high energetic cost of wrapping, reducing the thermodynamic stability of the nucleosome. Importantly, strong nucleosome positioning requires not only deformable DNA, but also an appropriate ∼10-bp periodic arrangement of A/T- and G/C-rich motifs that promotes rotational positioning on the histone surface. Because this periodic sequence pattern necessarily alternates highly deformable A/T-rich motifs with mechanically stiffer G/C-rich motifs, it precludes strong-positioning sequences from achieving the extreme flexibility of repetitive sequences such as polyAT. Consistent with this interplay between sequence patterning and DNA mechanics, integrated machine-learning models accurately predict genome-wide nucleosome positioning directly from DNA sequence and its intrinsic physical properties^88^. The convergence of genomic sequences and artificially optimised strong-positioning sequences within this intermediate regime indicates that robust nucleosome positioning is achieved by balancing DNA deformability with rotational sequence patterning, resulting in nucleosomes that are both thermodynamically stable and mechanically responsive. We hypothesize that such a balance preserves nucleosome thermodynamic stability while enabling regulated unwrapping in response to histone modifications and physiological forces.

An important consequence of this narrow mechanical regime is that DNA mechanics alone are unlikely to account for the large differences in nucleosome stability observed in cells. This mechanical regime should therefore be interpreted within the broader context of chromatin organisation *in vivo*, where nucleosome behaviour is governed not only by intrinsic DNA sequence but also by histone chemistry, chromatin remodelling activities, transcription factors, other chromatin-binding proteins, and the local chromatin environment. Indeed, our simulations reveal that even in the absence of additional factors, histone chemistry is the dominant tuning parameter for nucleosome stability, capable of exerting strong cooperative effects. This arises from the energetic balance between the cost of bending DNA to wrap it around the histone core versus the gain provided by histone–DNA electrostatic interactions. Altering histone charge through post-translational modifications produces the most pronounced effects. Lysine acetylation substantially weakens histone–DNA electrostatics and leads to large reductions in force barriers, particularly for the inner DNA turn. In contrast, histone variants induce more modest but consistent changes to nucleosome unwrapping. For example, CENP-A nucleosomes exhibit a pronounced reduction of the first unwrapping barrier, while H2A.Z reduces the second barrier, consistent with enhanced nucleosome breathing and prior experimental observations^67,89^.

At amino-acid resolution, our simulations reveal that wrapping of the DNA around the histone core is predominantly contributed by the arginines in the histone core (particularly in H3 and H2A) and the lysines in the flexible tails (especially in H3, H2B). Notably, tail–DNA contacts persist even as core–DNA contacts are lost, helping to stabilise partially unwrapped states. These trends align with experimental data on tail residence times^54^, truncation mutants^90^, and single-molecule pulling assays^41^.

When DNA mechanics are deliberately perturbed outside the ‘Goldilocks zone’ of intermediate flexibility, the interplay between DNA sequence and histone chemistry becomes even more pronounced. Histone modifications that weaken electrostatic interactions, such as tail acetylation, destabilise nucleosomes most severely when combined with mechanically rigid DNA (e.g., polyA), facilitating force-induced unwrapping. Conversely, pairing the same acetylated histones with more flexible DNA (e.g., polyAT) partially mitigates this destabilisation, restoring mechanical resistance and revealing a compensatory mechanism between DNA mechanics and histone chemistry. Analogous behaviour is observed for histone variants: while TP53 DNA destabilises nucleosomes assembled with canonical histones, this effect is modestly enhanced in H2A.Z-containing nucleosomes, indicating that H2A.Z can act cooperatively with the DNA sequence on nucleosome destabilisation.

It is important to distinguish between spontaneous nucleosomal breathing and force-induced unwrapping. Breathing involves short-lived, low-amplitude fluctuations of the entry–exit DNA triggered by thermal fluctuations, whereas unwrapping requires external forces, such as those applied in this work, experimentally^31^ or by ATP-dependent chromatin remodellers^91^. While histone tail acetylation is known to increase both breathing^24^ and susceptibility to unwrapping under force, as revealed here, enhanced breathing arising from DNA sequence alone does not necessarily imply reduced mechanical resistance for unwrapping. In such cases, the electrostatic landscape anchoring DNA to the histone core remains largely unchanged, and tail–DNA contacts are not inherently easier to disrupt under tension. More generally, factors that directly alter histone–DNA electrostatics or tail conformations are likely to play a decisive role in reshaping the unwrapping energy landscape of nucleosomes.

This study advances our understanding of how nucleosome physicochemical parameters encode both nucleosome stability and plasticity. We show that nucleosomes behave as mechanical systems in which DNA elastic deformation, histone–DNA electrostatics, and nucleosome geometry jointly shape the response to force. These physical principles have direct implications for understanding chromatin-mediated processes such as transcription, replication, and DNA repair, where nucleosomes are subjected to mechanical stresses. The force–response landscapes we present in this work reveal how DNA mechanics and histone chemistry set the intrinsic mechanical barriers that ATP-dependent chromatin remodellers^92^ must overcome.

Together, our results provide a mechanistic framework for how DNA sequence, nucleosome geometry, and histone chemistry cooperatively regulate nucleosome stability and accessibility. In genomic nucleosomes, DNA sequences occupy a narrow mechanical regime that supports stable yet plastic nucleosomes, while histone composition acts as a powerful regulatory knob that can strongly enhance or weaken stability when required. This interplay reveals how the extensive physicochemical diversity of nucleosomes intrinsically encodes a broad repertoire of DNA accessibility behaviours that can be differentially exploited by chromatin remodellers.

## Methods

### Chemically-specific chromatin coarse-grained model

Our chemically-specific coarse-grained model for chromatin^30^ meets two opposing requirements. The first is to achieve a high enough resolution to accurately represent proteins and DNA, capturing the effects of variations in DNA and amino acid sequences on nucleosome-nucleosome interactions, as well as the intrinsic nucleosome breathing motions that naturally arise from histone–DNA interactions and the DNA’s mechanical properties. The second is to minimise the degrees of freedom within chromatin, enabling efficient simulation of oligonucleosome systems. Although computationally demanding, describing proteins at one bead per amino acid and DNA at one ellipsoid per base pair is essential to achieving these goals. This resolution preserves the full chemical diversity of a heterogeneous chromatin array, allowing us to map the varied physicochemical properties of the distinct amino acids and nucleobases that constitute different nucleosomes within chromatin. This level of detail also allows for the consideration of point mutations in amino acids, if needed, to investigate the role of specific histone residues in chromatin organisation, assess the binding of additional proteins to specific chromatin sites, and analyse the contributions of individual amino acids to nucleosome unwrapping. Moreover, while histone core proteins are largely *α*-helical and exhibit minimal structural fluctuations in crystallographic studies^1,37^ and atomistic MD simulations^93^, histone tails are mostly disordered and highly flexible. A resolution of one bead per residue is the coarsest that can still accurately describe the detailed topology of the histone globular domains, the flexibility of the histone tails, and potential disorder-to-order transitions triggered by post-translational modifications^6^. Accordingly, the histone globular domain is enforced using a Gaussian elastic Network Model (GNM) to preserve the integrity of the histone folding, while the histone tails are modeled as fully flexible polymer chains capable of undergoing large conformational fluctuations. For DNA, a resolution of one bead per base pair is the coarsest that can fully capture the influence of DNA sequence on its mechanical properties, such as twist, roll, and tilt. Collectively, our model reduces the number of particles in a 5 kb chromatin region, or a system of approximately 25 nucleosomes, from about 0.5 million atoms (plus solvent) to only around 25,000 beads. Further details of this model are provided below.

Another key feature of our model is its consideration of sequence-dependent DNA mechanical properties, implemented through the Rigid Base Pair (RBP)^74,94–97^ model with added phosphate charges. The RBP model represents each DNA base-pair step with a single ellipsoid, depicting DNA conformational changes via harmonic deformations of six helical parameters (three angles: twist, roll, and tilt, and three distances: slide, shift, and rise), which account for the relative orientations and positions of neighbouring base-pair planes. The DNA mechanical potential energy is calculated by summing the harmonic distortions of equilibrium base-pair step geometries. We used parameters from the Orozco group^74,97^, derived from MD atomistic simulations—specifically, equilibrium values fitted with Gaussian functions to the distributions of helical parameters, and elastic force constants obtained by inverting the covariance matrix in helical space. In practice, our model represents single base pairs, both within nucleosomal and linker DNA, with one coarse-grained bead defined by a position vector, **r**, and an orientation quaternion, *<u>q</u>*. We add two virtual charge sites to each DNA ellipsoid (one per phosphate, approximating the shape of the DNA phosphate backbone) to account for the crucial electrostatic interactions that drive chromatin self-organisation. This RBP plus charged virtual sites model is implemented in LAMMPS^98^, with an ellipsoid defined by two-point particles of negligible but non-zero mass. While the combined three-particle base-pair bead is treated as a single rigid body for dynamics, each component contributes to the calculation of potential and forces.

Beyond excluded volume and hydrophobic non-bonded interactions, we incorporate electrostatic interactions between the charged beads using a Debye-Hückel potential, while omitting non-bonded interactions between directly bonded beads. The binding of nucleosomal DNA to the histone protein core is achieved through these protein–DNA electrostatic and hydrophobic interactions, resulting in nucleosomal DNA that wraps approximately 1.7 times around the histone core and exhibits spontaneous unwrapping and re-wrapping. The force-induced unwrapping behaviour of these nucleosomes quantitatively agrees with experimental data at single base-pair resolution^31,32,39–42,44^. For modelling nucleosomes artificially constrained to be ‘non-breathing’, we describe the histone core together with the bound nucleosomal DNA as a single Gaussian Network Model (GNM), using the same 7.5 Å threshold and bond parameters. This approach constrains the nucleosomal DNA to remain permanently bound to the histone core, preventing both nucleosome breathing and sliding.

In summary, our chemically-specific coarse-grained model of chromatin preserves the atomistic shape and size of the nucleosome core, the length and flexibility of the histone tails, the explicit charges and hydrophobic nature of all amino acids within the histone protein (including those in the critical acid patch region^1^), the sequence-dependent mechanical properties of DNA, and the thermal breathing motions of nucleosomes. A comprehensive list of all parameters used in our chemically-specific model, along with the chosen values and their justifications, is provided in Ref.^30^

### Nucleosome pulling simulations

The initial structure for our simulations is modeled from a high-resolution crystal structure of a nucleosome (named 1KX5 from now on for its PDB accession code^37^). The associated DNA is formed by 211-bp construct: the 147-bp nucleosomal DNA sequence from 1KX5 flanked symmetrically by two 32-bp linker DNA arms. These linker sequences were taken from the 1ZBB tetranucleosome crystal structure (Chain J, the full sequence: nucleotides 159–347; GenBank accession 1ZBB_J)^38^. Unless specified, we use this sequence by default. For nucleosomes containing the histone variants H2A.Z and CENP-A, the initial structure is based on the coordinates deposited in the Protein Data Bank with the accession codes 1F66^56^ and 6TEM^65^, respectively. For the H3.3 variant, the mutation is introduced into the 1KX5 structure. Rigid DNA base pairs were fitted to the all-atom structure using the software x3DNA 3.0^99^ (position vector and an orientation matrix were obtained) and then mutated by changing the base pairs parameters (see SI for the full list of sequences). PTMs were modeled by setting the charge of acetyl-lysine to 0 and increasing the sigma accordingly to addition of an acetyl group. All the initial structures consist of the histone octamer fully wrapped by 211-bp DNA. Each system was relaxed for 2000000 steps (80 ns). An Umbrella Sampling protocol was used to calculate the free energy curve along the extension. To generate initial configurations for each window, a constant velocity Steered MD was performed, from an initial distance between DNA strands of 25 Å extension up until an extension of 750 Å. During the steered MD, a spring constant of 0.01 kcal/mol/Å^2^ was used to add a harmonic restraint, whereas a velocity of 9.0 x 10-6 Å/fs was used to mimic the force-spectroscopy experiments setup. The extension range was then divided into 60 evenly spaced windows. Each window was simulated for 40 ns using a fixed harmonic biasing potential at the corresponding extension, with a spring constant of 0.025 kcal/mol/ Å^2^ using the COLVARS package from LAMMPS^100^ (Convergence test: Fig. S1). For each simulation, we performed five replicas, ensuring independence by setting up the procedure with different random seeds.

The combined data of each pulling experiment was then used to compute the Potential of Mean Force using the Weighted Histogram Analysis Method (WHAM)^34^ (version 2.0.9) (Supplementary Table 5). Each of our simulations was performed over several independent replicas. Furthermore, since we perform each simulation via umbrella sampling, each simulation point is independently converged and, therefore, independent from the previous point of the curve.

#### Quantification of unwrapping free energies

The equilibrium free-energy cost of nucleosome unwrapping was defined as ΔΔ*G* = Δ*G*(*x*_unwrapped_) − Δ*G*(*x*_wrapped_), where Δ*G*(*x*) denotes the PMF value at extension *x*, with 25 Å corresponding to the fully wrapped state and 620 Å corresponding to the fully unwrapped state. All reported ΔΔ*G* values therefore represent the equilibrium free-energy difference between the wrapped and unwrapped configurations.

To enable direct comparison between modified nucleosomes and the canonical 1KX5 system, absolute differences in unwrapping free energy were computed as Δ(ΔΔ*G*) = ΔΔ*G*^mod^ − ΔΔ*G*^1KX5^. Uncertainties in ΔΔ*G* correspond to the standard error of the mean (SEM) over five independent replicas. The uncertainty in the absolute free-energy difference was obtained by standard error propagation assuming independent simulations,

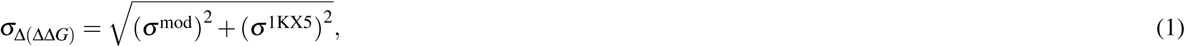

where *σ* ^mod^ and *σ* ^1KX5^ denote the SEM values of the modified and canonical systems, respectively.

#### Pulling Force Calculation

Force–extension curves were reconstructed by numerical differentiation of the individual PMF calculation with respect to the reaction coordinate, using a central finite difference scheme. Force profiles were averaged over the five independent replicas for each condition. Two characteristic unwrapping events were defined: F1 (first peak) and F2 (second peak), corresponding to force maxima. For each peak, the peak magnitude and its coordinate were extracted from each replica and averaged to obtain condition–level means and standard error of the mean (s.e.m). For each condition, the difference in the F2 peak position relative to the reference state (1KX5) was defined as Δ*X*_F2_. Errors inΔ*X*_F2_ were obtained from the s.e.m of the underlying coordinate distributions.

To quantify changes in mechanical stability relative to the canonical nucleosome (1KX5), percentage differences in the force barriers were computed for each modified condition as

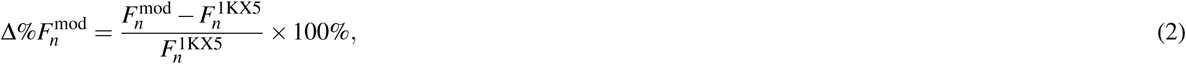

with *n* = 1, 2 corresponding to the outer and inner DNA turns, respectively. The associated uncertainty was computed using standard error propagation:

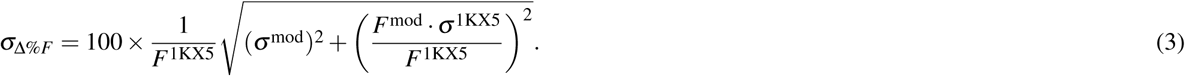

For each condition, the difference in the position of the second force peak relative to the reference state (1KX5) was defined as Δ*X*_F2_. Errors in Δ*X*_F2_ were obtained from the s.e.m. of the underlying coordinate distributions.

#### Angle Calculation

The tilt/reorientation angle was defined to quantify the nucleosome core reorientation relative to the pulling direction along the extension. The angle was computed from the atomic coordinates using a custom VMD^101^ script. For each trajectory frame, four sets of atoms were selected: (i) a dyad–region base–pair stack, (ii) an antidyad–region stack positioned approximately opposite to the dyad, and (iii–iv) two segments on the left and right DNA gyres. The geometric centre of each selection was computed. All atomic coordinates in the frame were then translated such that the antidyad centre coincided with the origin. A dyad–antidyad axis vector was defined as the displacement from the dyad centre to the antidyad centre, and a second vector was defined between the centres of the left and right gyres. The normal vector of the nucleosome disc was then obtained as the negative cross product of these two in–plane vectors and normalized to unit length. The reorientation angle *θ* was defined as *θ* = arccos **n̂** · **k̂**, where **n̂** is the normalized disc normal and **k̂** = (0, 0, 1) is the pulling axis, and therefore ranges from 0 to 180 degrees. The angle was computed for every frame and recorded as a time series *θ* (*t*). It measures the instantaneous orientation of the nucleosome relative to the pulling direction, rather than a signed rotational coordinate. We note that we use this definition of the tilt/reorientation angle as it has been used in several important works on nucleosome unwrapping, including those by Schiessel and co-workers, Arya and co-workers and Thirumalai and co-workers^36,46,47^ to characterise the nucleosome reorientation transition proposed to accompany force-induced unwrapping. Using a consistent definition allows us to compare our predictions to those of others.

#### Contact Analysis

Histone–DNA contacts along the reaction coordinate were quantified by computing distance-based contact maps for each umbrella–sampling window and replica. Contacts were defined exclusively between DNA phosphate groups and lysine or arginine residues, using a 7.5 Å cutoff. The contact frequency for each base pair was visualized as a heatmap as a function of end-to-end extension.

Electrostatic interactions between basic residues and DNA were evaluated using a Debye–Hückel formalism. Distances between lysine or arginine residues and DNA phosphates were extracted from 60 windows spanning 25–725 Å of nucleosome extension, with five replicas per window; only residue–phosphate pairs within 7.5 Å were considered. The interaction energy for each pair was computed using the Debye–Hückel potential (SI eq. 3) with *q*_1_ = +1, *q*_2_ = −1, relative permittivity *ε_r_* = 80, and Debye length *λ_D_* = 8 Å, corresponding to 150 mM monovalent salt. Total electrostatic contributions were computed per replica and averaged across replicas for each window.

#### DNA Elastic Deformation Energy Calculation

The elastic deformation energy was calculated as 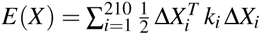 where *X* denotes the six–dimensional vector of helical parameters (Shift, Slide, Rise, Tilt, Roll, and Twist), and Δ*X_i_* is the deviation vector for the base–pair step *i*. For each step, Δ*X_i_*the helical parameters of the equilibrated, fully wrapped nucleosome configuration immediately prior to the onset of the pulling protocol were obtained, thereby capturing the deformation imposed on DNA while still fully engaged with the histone core. The reference equilibrium helical values correspond to naked DNA extracted from the ABC microsecond simulations^74^. The associated 6 × 6 stiffness matrices *k_i_* are sequence–dependent tensors derived from the same ABC trajectories, which were also used to construct the RBP model.

The relationships among DNA deformation energy, ΔΔ*G*, and the position of the second unwrapping barrier were quantified from umbrella sampling PMFs. The free energy was evaluated at an extension of 620 Å and referenced to the fully wrapped nucleosome state, yielding a relative free energy difference. This procedure was applied independently to each replica, and per-sequence mean values and standard errors of the mean (s.e.m.) were computed as shown above. Linear least-squares fits and Pearson correlation coefficients were computed directly from the plotted data. Error bars represent standard errors across five replicas for each observable, and axes are reported in units of kcal mol^−1^ (ΔΔ*G*), kcal mol^−1^ (energy), and Å (coordinate).

### Software used

Simulations were performed using LAMMPS^98^ (version 3rd March 2020) with our custom code ^30^. We used the program 3DNA^99^ (version 2.3) within our model building methods. All data analysis was done using Python (version 3.8.5) with NumPy (version 1.19.2) and SciPy (version 1.5.2). All data were plotted using Matplotlib (version 3.3.2). The reorientation angle was calculated using VMD^101^. Images were rendered using Tachyon render of the Open Visualization Tool (OVITO Pro) software^102^ (version 3.10.5). We used the Weighted Histogram Analysis Method (WHAM) program^34^ (version 2.0.9) to calculate PMFs. All simulations were executed using the computational resources of the Cambridge Service for Data Driven Discovery (CSD3) and the Archer2 High Performance Computing facility.

## Code Availability

The authors are delighted to share the computational implementation of their models with the community. All the necessary files can be found at: https://github.com/CollepardoLab/CollepardoLab_Chromatin_Model and https://doi.org/10.6084/m9.figshare.13663685.v1. Please use them freely and remember to cite this paper and LAMMPS (http://lammps.sandia.gov)^98^. The authors are happy to answer any questions and comments by email and welcome contributions for any updates. For visualization of the free energy curves, you can visit https://janhuemar.github.io/PullingNucleosomes/

## Acknowledgements

This work was funded by the UK Research and Innovation (UKRI) Engineering and Physical Sciences Research Council (EP-SRC) under the UK Government’s guarantee scheme (grant EP/Z002028/1 awarded to RCG). MJM would like to acknowledge the Winton Programme for Physics of Sustainability for doctoral funding. JIPL acknowledges the Gates Cambridge Trust for doctoral funding. JH is supported by the Herchel Smith Postdoctoral Fellowship Fund. We acknowledge EuroHPC Joint Undertaking for awarding access to MareNostrum5 at Barcelona Supercomputing Center (BSC), Spain [EHPC-REG-2025R01-166]. This project also utilised time on HPC resources granted by the UK High-End Computing Consortium for Biomolecular Simulation, HECBioSim (http://hecbiosim.ac.uk), supported by EPSRC (grant no. EP/X035603/1) to RCG.

## Author contributions statement

M.J.M., J.H., and R.C.G. conceived the project. M.J.M., J.H., and J.I.P.L. designed the research. S.F. and M.J.M. developed the computational framework and software. M.J.M. conceived and developed the theoretical models. J.I.P.L. performed all simulations. M.J.M., J.I.P.L. and J.H. analysed the data and prepared the visualizations. J.H. and R.C.G. supervised the research, and R.C.G. acquired funding. J.H. and R.C.G. wrote the original draft of the manuscript. All authors contributed to reviewing and editing the manuscript and approved the final version.

