## Supplementary Figure for "A Goldilocks zone of DNA flexibility defines stable yet plastic nucleosomes, tuned by histone chemistry"

**Supplementary Tables** **14**

**Supplementary References** **17**

#### Supplementary Figures

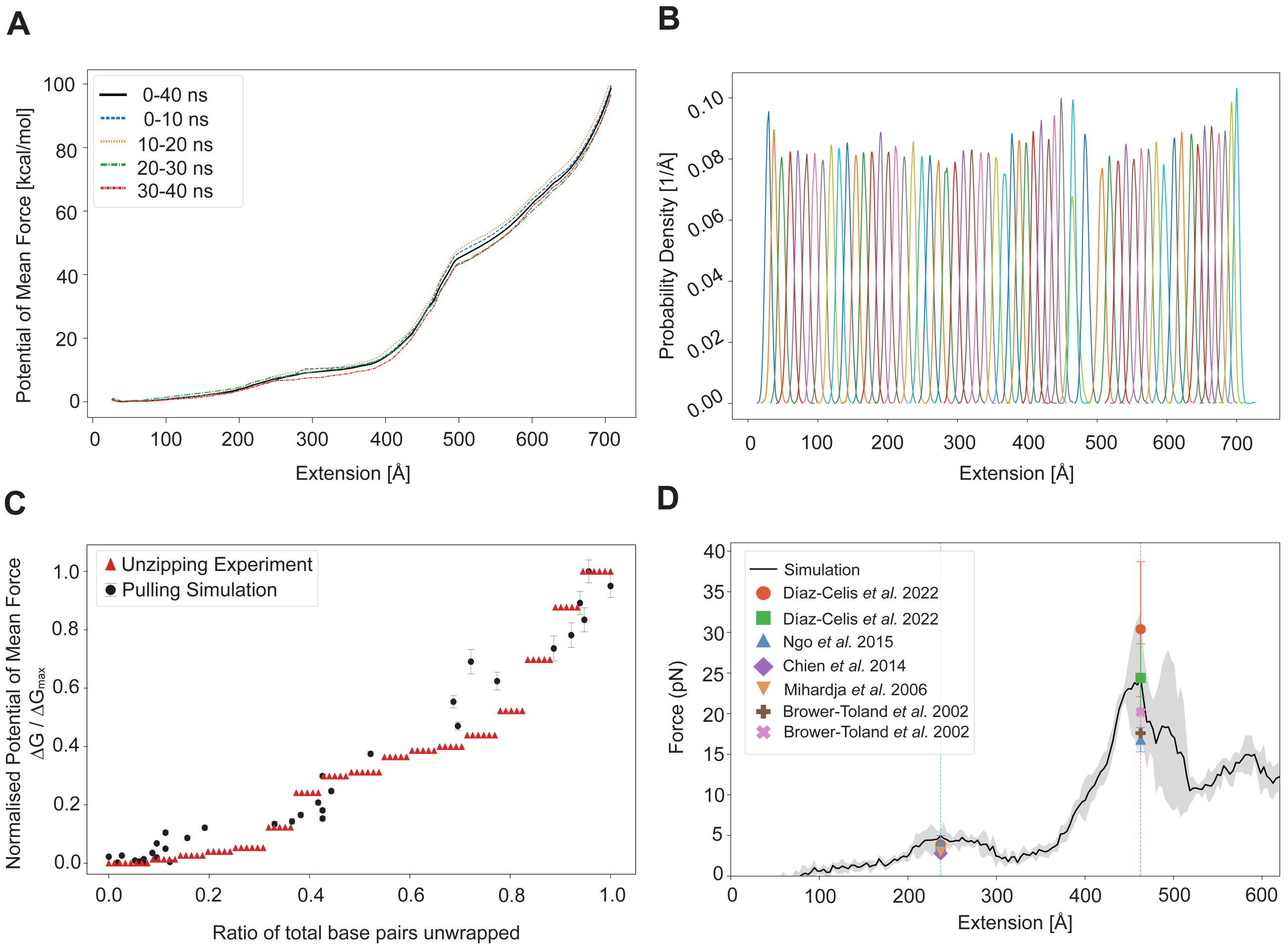

**Supplementary Figure 1. Convergence analysis and Comparison with Experiments. A.** Block analysis of the umbrella sampling PMF calculation obtained with WHAM. The PMF was recomputed using successive 10 ns blocks (0–10 ns, 10–20 ns, 20–30 ns, 30–40 ns) and the full 0–40 ns dataset, showing convergence of the free-energy profile and the absence of drift over the sampled time intervals. **B.** Normalised probability densities for all umbrella windows. The substantial overlap between neighbouring histograms confirms that the chosen window spacing and biasing potentials provide sufficient sampling continuity for reliable WHAM reconstruction of the PMF. See Table 5 for the histogram overlap of all conditions**C.** Normalised potential of mean force for DNA unwrapping from a nucleosome on the 601 positioning sequence from unzipping experiment [1] in red compared with our simulation values for the same sequence in black. **D.** Force–extension unwrapping of a single nucleosome in a 601 sequence, obtained by taking the numerical derivative of the potential of mean force values from umbrella sampling simulations. Markers are placed to represent the experimental values reported for the first unwrapping barrier (F1, green dashed line) and the second unwrapping barrier (F2, orange dashed line). See Table 1 for the exact values, experimental conditions and references.

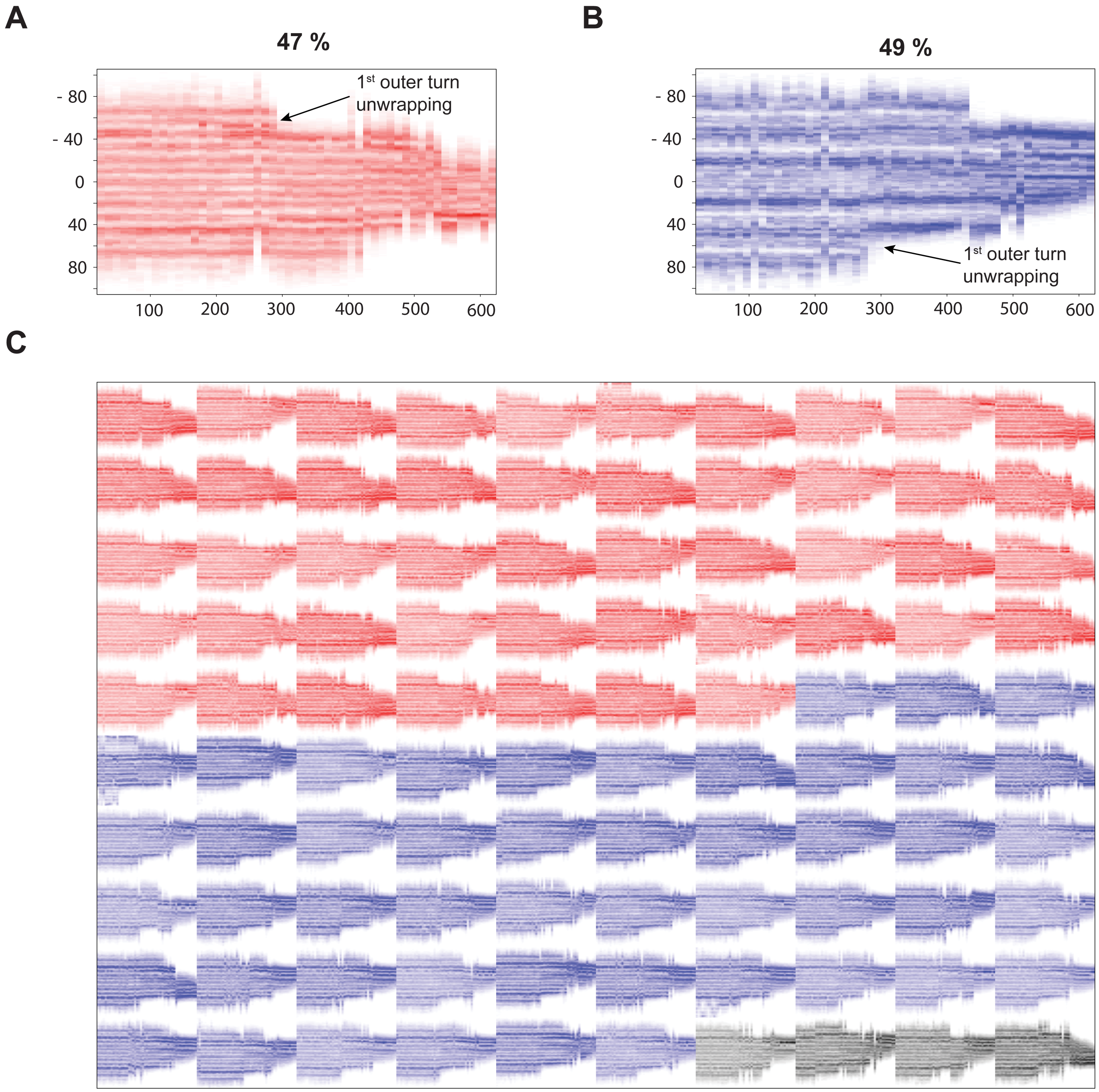

**Supplementary Figure 2. Asymmetric first-turn unwrapping. A.** DNA–protein contact heatmap (base-pair resolution) as a function of end-to-end extension for trajectories in which the *right* DNA arm unwraps first (47% of all simulations). **B.** Heatmap example for trajectories in which the *left* DNA arm unwraps first (49%). **C.** Complete set of 100 individual contact heatmaps used for classification. The top block corresponds to right-first unwrapping (red), the middle block to left-first unwrapping (blue), and the bottom block shows the four trajectories that could not be unambiguously assigned.

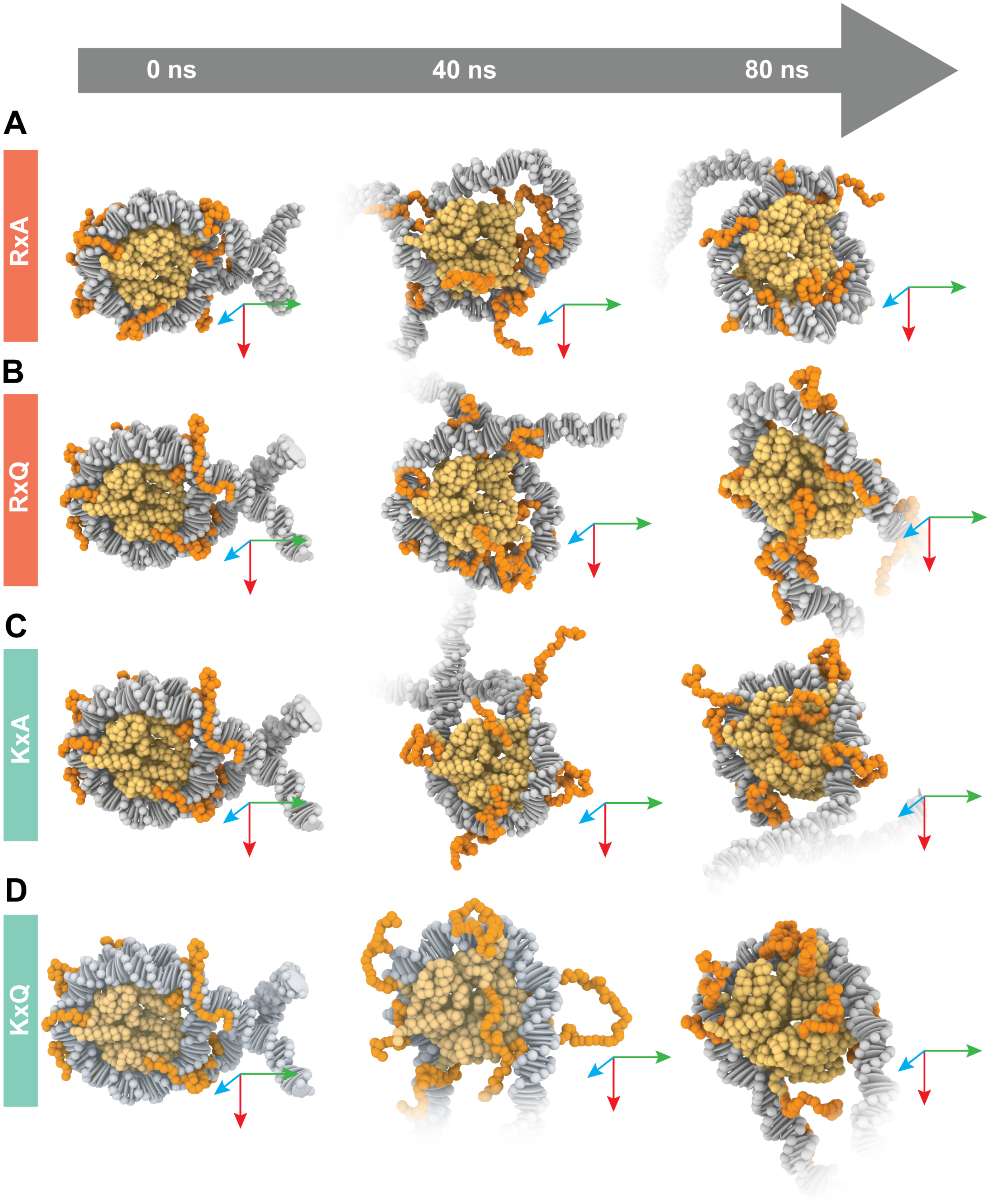

**Supplementary Figure 3. Spontaneous unwrapping simulations. A.** Snapshots of arginine-to-alanine mutants. **B.** Snapshots of arginine-to-glutamine mutants. **C.** Snapshots of lysine-to-alanine mutants. **D.** Snapshots of lysine-to-glutamine mutants. All the mentioned residues were replaced, followed by an unbiased molecular dynamics simulation. The nucleosomes were completely unwrapped within the first 80 ns of the simulation without the application of external forces in all replicas.

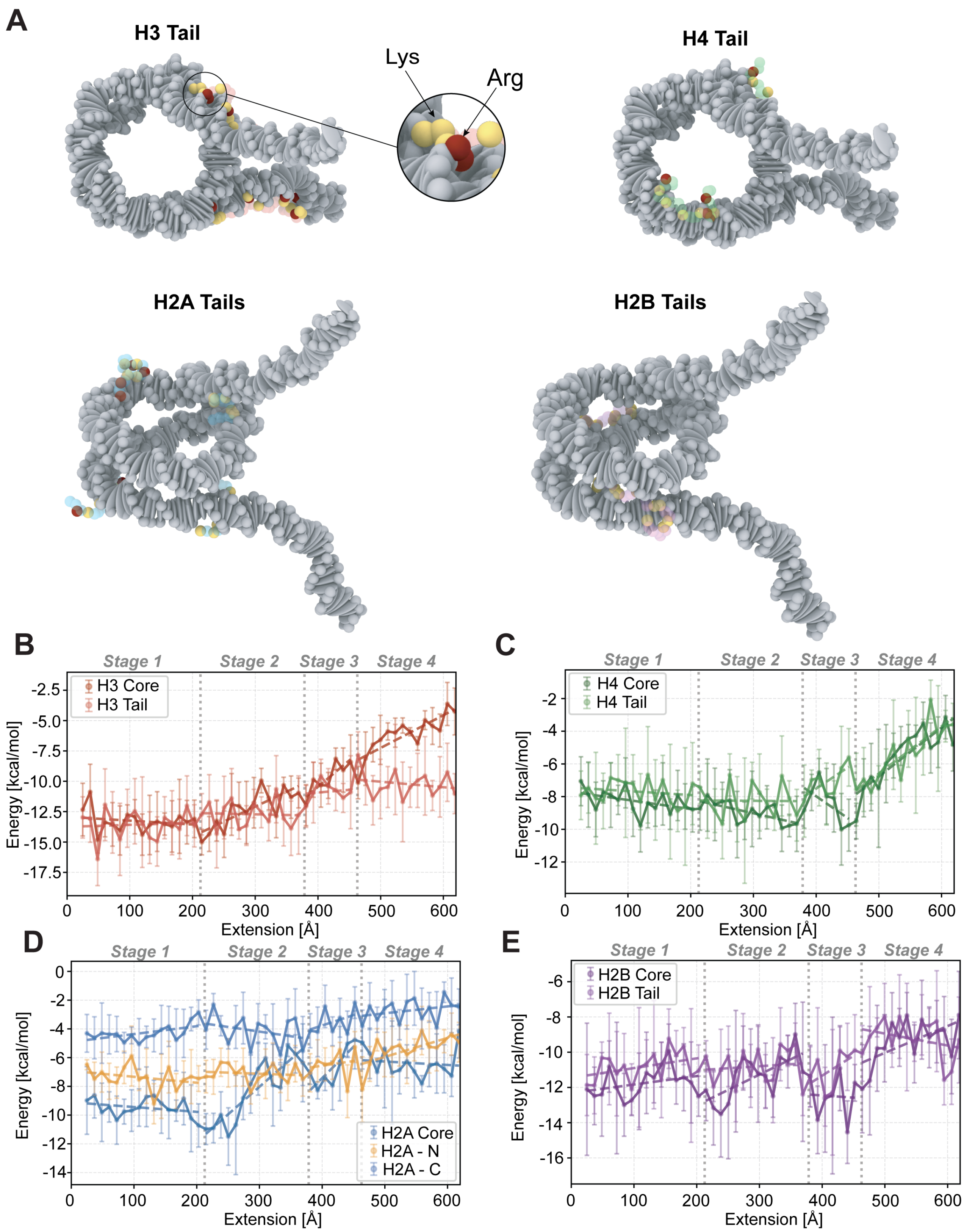

**Supplementary Figure 4. Time-resolved lysine and arginine contributions by tail during nucleosome unwrapping. A.** Models of nucleosomal DNA (grey) and histone tails (transparent) showing the positions of lysines (solid beads in yellow) and arginines (solid beads in dark red). Electrostatic energy calculated from the arginine and lysine protein–DNA contacts using a Debye–Hückel approach, as a function of the end-to-end extension coordinate: **B.** for H3, **C.** H4, **D.** H2A, and **E.** H2B.

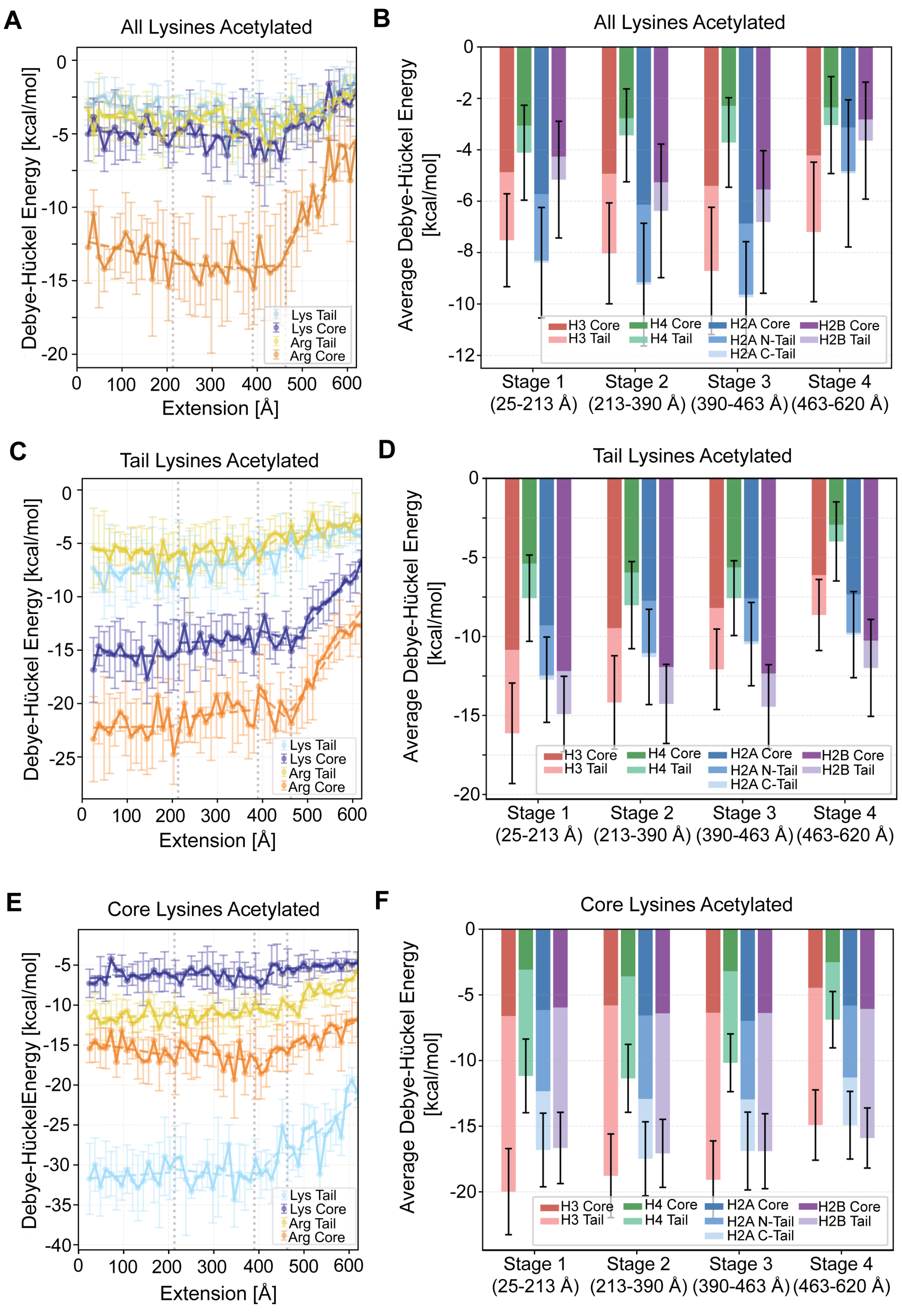

**Supplementary Figure 5. Differential impact of acetylation in the histone tails and the histone core. A, C, E.** Debye–Hückel electrostatic energy between positively charged histone residues (lysines and arginines) and DNA as a function of the end-to-end extension coordinate, shown separately for nucleosomes in which (A) all lysines are acetylated, (C) only tail lysines are acetylated, and (E) only core lysines are acetylated. Energies are decomposed into contributions from lysine tails, lysine cores, arginine tails, and arginine cores. **B, D, F.** Average Debye–Hückel energy of each histone segment across the four unwrapping stages (Region 1: 25–213 Å, Region 2: 213–390 Å, Region 3: 390–463 Å, Region 4: 463–620 Å), for the same acetylation schemes as in A, C, and E. Energies are resolved by histone type (H3, H4, H2A, H2B) and by core and tail contributions.

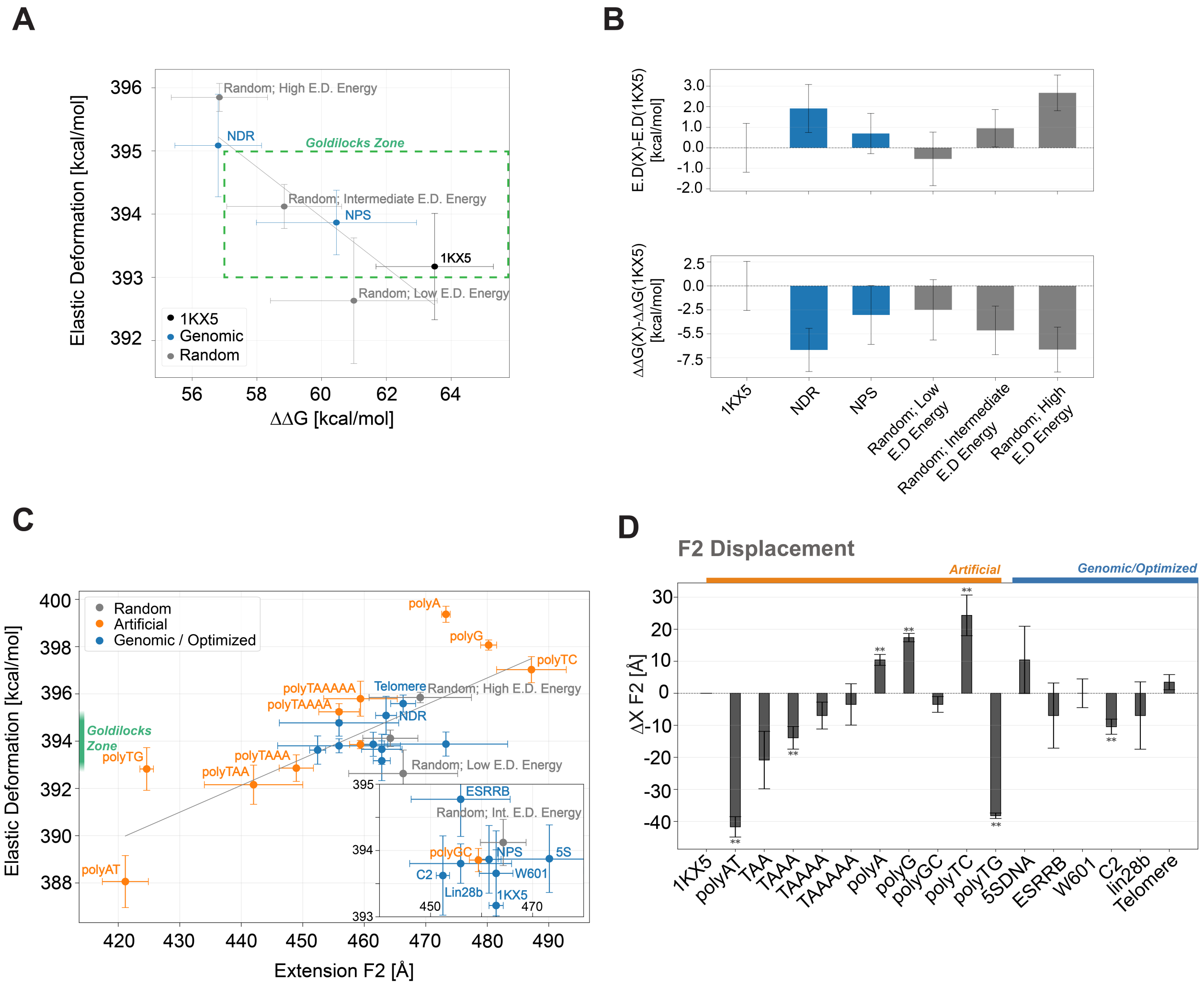

**Supplementary Figure 6. Effects of DNA elastic deformation energy in nucleosome unwrapping. A.** $\Delta\Delta G_{620}$ of nucleosome unwrapping plotted against the elastic deformation energy of wrapped DNA. Lower $\Delta\Delta G_{620}$ values indicate greater mechanical plasticity and easier unwrapping under force, whereas higher elastic deformation energy is associated with reduced nucleosome stability. The green box represents the "Goldilocks zone". The 1KX5 sequence is shown in black as a reference. Yeast NDR sequences from TSS-flanking regions show higher elastic deformation energy and lower $\Delta\Delta G_{620}$ than yeast nucleosome-positioning sequences (NPS). Random sequences are shown in gray and illustrate that variation in DNA elastic deformation energy partly encodes differences in nucleosome stability. However, even random sequences with lower elastic deformation energy than the 1KX5 do not achieve the stability of this sequence, supporting that other sequence-dependent features, such as TA periodicity, contribute to nucleosome stability. **B.** Differences of elastic deformation (top) and the $\Delta\Delta G_{620}$ (bottom) with respect to the 1KX5 condition. **C.** Correlation between $\Delta X_{\mathrm{F2}}$ and elastic deformation energy. Artificial poly-sequences are shown in orange, genomic nucleosome-forming optimised sequences are shown in blue and random sequences in grey. The correlation presents an $R^2$ of 0.60. The error bars represent the s.e.m. **D.** Shift in the position of the second unwrapping barrier, $\Delta X_{\mathrm{F2}}$, for all sequences. Positive values indicate delayed unwrapping; negative values indicate earlier unwrapping.

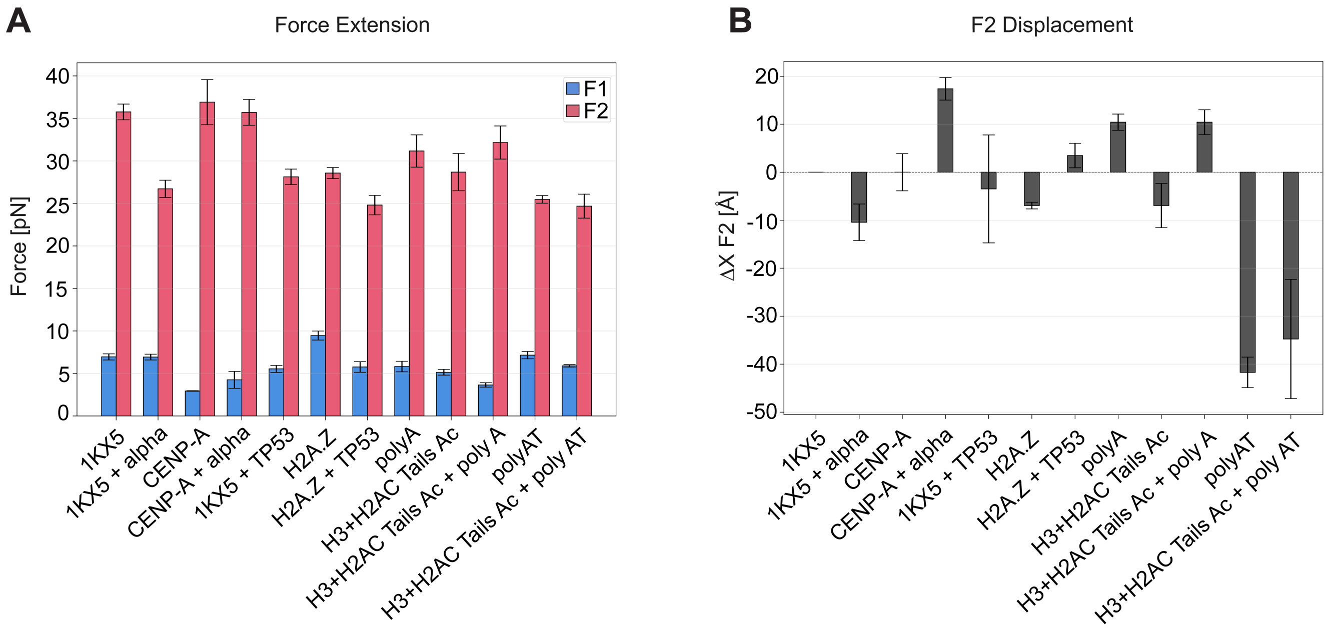

**Supplementary Figure 7. Concerted changes in DNA sequence and histones modulate nucleosome unwrapping.** Forces associated with the first (F1) and second (F2) unwrapping barriers for all combined conditions studied, including histone variants (CENP-A, H2A.Z), natural DNA sequences (TP53), artificial repetitive sequences (polyA, polyAT), and systems containing both H3+H2A C-terminal tail acetylation and sequence perturbations. Bars report mean values over five replicas, with error bars indicating the standard error.

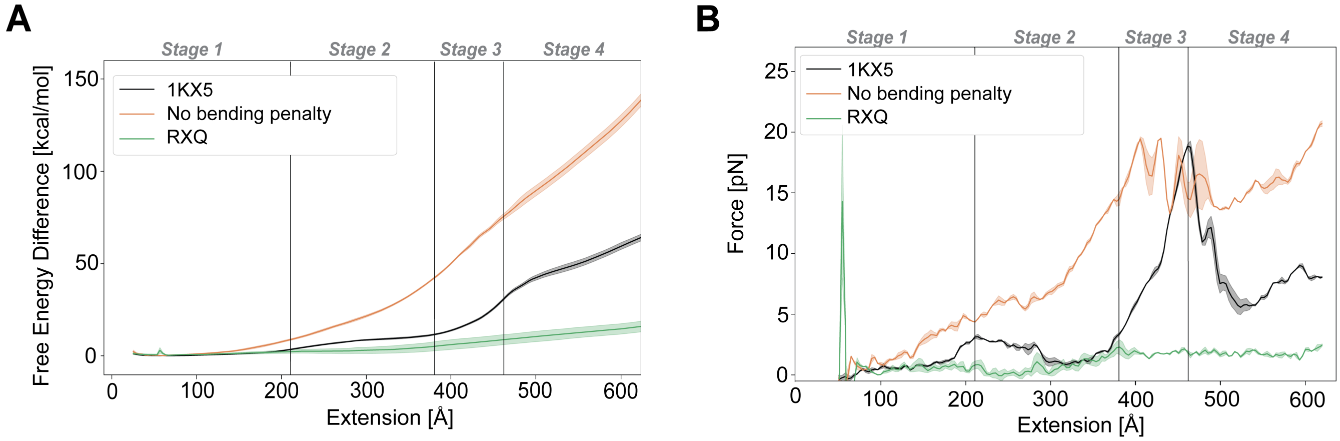

**Supplementary Figure 8. DNA flexibility and histone charge modulate nucleosome unwrapping. A.** Free-energy difference as a function of end-to-end extension for a 1KX5 nucleosome containing 211 base pairs of DNA: the reference system, a system with no energetic bending penalty for DNA (tilt ($\tau_n$) and roll ($\rho_n$) coefficients in $K_n$ of the $E_{RBP}$ expression (1) set to zero), and a system in which all arginines are mutated to glutamines. **B.** Force–extension unwrapping curve obtained by taking the numerical derivative of panel A.

### Supplementary Notes

#### Model Definition

##### Energy Function

In our high-resolution coarse-grained model, each DNA base pair is represented with an ellipsoid and two virtual negative charges to represent the DNA phosphates. The DNA configurational potential is modelled using a Rigid Base Pair (RBP) approach, where inter-base-pair step deformations are sequence-dependent and effects are captured through six helical parameters: twist, roll, tilt, slide, shift, and rise. That is, the configurational potential energy of the DNA is computed as the sum of harmonic distortions from these helical parameters, with equilibrium values and stiffness values derived from atomistic simulation datasets [2, 3]:

$$E_{\mathrm{RBP}} = \sum_{n} \frac{1}{2} \Delta\phi_n K_n \Delta\phi_n^{\mathrm{T}} \tag{1}$$

Here, we describe a harmonic potential for the bonded interactions, where $\Delta\phi_n$ denotes the deviation of the six helical parameters from their sequence-dependent equilibrium values and $(D_{x_n}, D_{y_n}, D_{z_n}, \tau_n, \rho_n, \omega_n)$ , $K_n$ is the corresponding $6 \times 6$ stiffness matrix for the given base pair. $\Delta\phi_n^{\mathrm{T}}$ denotes the transposed matrix of $\Delta\phi_n$

As mentioned above, each amino acid is simplified as a spherical particle that represents the key biophysical properties of interest, including net charge, hydrophobic character, and steric exclusion. Beads are spatially positioned at the positions of the $C_\alpha$ atoms in the all-atom structure. Bead sizes are derived from experimental van der Waals volumes, assuming a spherical approximation [4]. Histone tails are treated as fully flexible polymers with harmonic bonded potentials with no penalty for bending.

$$E_{\mathrm{Bonds}} = \frac{1}{2} k (r - r_0)^2 \tag{2}$$

The bonded interactions are defined as a harmonic potential where $k$ is the spring constant (10 kcal/mol/Å$^2$), $r$ is the current bond length and $r_0$ is the bond length in the equilibrium.

To enforce the secondary structure of the globular domain of histones, a Gaussian elastic network model (GNM) is employed [5, 6]. This approach compensates for the inherent limitations of coarse-grained models in preserving secondary structure.

Non-bonded interactions among charged particles (Phosphate–phosphate, amino acid–amino acid, and phosphate–amino acid) are modelled with a Debye-Hückel potential, where $q_i$ and $q_j$ denote the charges of the interacting particles, $r$ is the distance between them, $\epsilon_0$ is the vacuum permittivity, $\epsilon_r$ the relative permittivity, and $\kappa$ is the inverse Debye length, related to the ionic strength of the system:

$$E_{\mathrm{Electrostatic}} = \frac{q_i q_j}{4\pi\epsilon_0\epsilon_r r} e^{-\kappa r} \tag{3}$$

This mean-field formulation allows us to approximate the screening effects of monovalent ions in solution without including them explicitly. We can modulate ionic screening by adjusting the Debye length $\lambda_D$:

$$\kappa^{-1} = \lambda_D = \sqrt{\frac{\epsilon_0 \epsilon_r k_B T}{2 \times 10^3 N_A e^2 c}} \tag{4}$$

Here, $k_B$ the Boltzmann constant, $T$ the absolute temperature, $N_A$ the Avogadro constant, $e$ the elementary charge, $c$ and the monovalent salt concentration in mol/L.

For non-ionic non-bonded interactions among any interacting pair, we use a truncated and shifted Lennard-Jones potential, with pair-specific parameters taken from the Kim-Hummer model [4]. These interactions are implemented using a truncated and shifted Lennard–Jones potential:

$$E_{\mathrm{KH}} = \begin{cases} E_{\mathrm{LJ}} + (1 - \lambda)\epsilon, & \text{if } r \leq 2^{1/6}\sigma, \\ \lambda E_{\mathrm{LJ}}, & \text{else,} \end{cases} \tag{5}$$

$$E_{\mathrm{LJ}} = 4\epsilon \left[ \left( \frac{\sigma}{r} \right)^{12} - \left( \frac{\sigma}{r} \right)^{6} \right] \tag{6}$$

Here, $\lambda$ quantifies hydrophobic strength, $\epsilon$ defines interaction depth, $\sigma$ is the effective contact distance, and $r$ is the inter-bead separation. Interactions between DNA and histone residues encompass both electrostatics and short-range forces, enabling realistic dynamics such as DNA wrapping, unwrapping, and repositioning around the histone core. These interactions were parameterized by fitting RDF for phosphate-amino acid pairs from atomistic MD simulations [7]. In simulations designed to inhibit breathing motions, additional GNM restraints are introduced between histone residues and nearby DNA virtual charges (within 7.5 Å), effectively restricting DNA mobility and mimicking a more static nucleosomal configuration.

The total energy function of our high-resolution coarse-grained chromatin model is given by:

$$E = \sum_{\text{Protein bonds}} E_{\text{Bonds}} + \sum_{\text{DNA bonds}} E_{\text{RBP}} + \sum_{i} \sum_{j<i} E_{\text{Electrostatic}} + \sum_{i} \sum_{j<i} E_{\text{KH}} \tag{7}$$

#### Additional Algorithms

##### Quaternion to Rotation Matrix

A unit quaternion

$$q = (q_w, q_x, q_y, q_z), \tag{8}$$

satisfies

$$|q| = \sqrt{q_w^2 + q_x^2 + q_y^2 + q_z^2} = 1. \tag{9}$$

It can be transformed into a corresponding $3 \times 3$ rotation matrix through the standard quaternion–matrix relation:

$$A = \begin{pmatrix} q_w^2 + q_x^2 - q_y^2 - q_z^2 & 2(q_x q_y - q_w q_z) & 2(q_x q_z + q_w q_y) \\ 2(q_x q_y + q_w q_z) & q_w^2 - q_x^2 + q_y^2 - q_z^2 & 2(q_y q_z - q_w q_x) \\ 2(q_x q_z - q_w q_y) & 2(q_y q_z + q_w q_x) & q_w^2 - q_x^2 - q_y^2 + q_z^2 \end{pmatrix}. \tag{10}$$

##### Computation of Helical Parameters

To extract helical parameters for the RBP representation, we follow the SCHNAaP protocol [8]. Consider two base pairs (represented as DNA ellipsoids) located at positions $\mathbf{r}_1$ and $\mathbf{r}_2$ with associated orientation quaternions $q_1$ and $q_2$. Our goal is to determine the set of geometric descriptors

$$\boldsymbol{\phi} = (D_x, D_y, D_z, \tau, \rho, \Omega),$$

which characterise their relative displacement and orientation. The procedure is as follows:

1. Convert quaternions $q_1$ and $q_2$ into direction–cosine matrices $T_1$ and $T_2$, whose columns correspond to the local $\hat{x}$, $\hat{y}$, and $\hat{z}$ axes.
2. Compute the roll–tilt bending magnitude,
$$\Gamma = \cos^{-1}(\hat{z}_1 \cdot \hat{z}_2). \tag{11}$$
3. Determine the roll–tilt axis,
$$\mathbf{rt} = \hat{z}_1 \times \hat{z}_2. \tag{12}$$
4. Rotate $T_1$ and $T_2$ by $\pm\Gamma/2$ around $\mathbf{rt}$:
$$T_1' = R(\mathbf{rt}, +\Gamma/2)\, T_1, \tag{13}$$
$$T_2' = R(\mathbf{rt}, -\Gamma/2)\, T_2, \tag{14}$$
where $R(\mathbf{a}, \theta)$ denotes the orthogonal rotation matrix corresponding to a rotation of angle $\theta$ about axis $\mathbf{a}$.

5. Define the mid–step frame as the average of the transformed frames,

$$T = \frac{1}{2}(T_1' + T_2'). \tag{15}$$

6. Evaluate the twist angle $\Omega$ from the transformed $\hat{y}$ vectors (second columns of $T_1'$ and $T_2'$):

$$\Omega = \cos^{-1}(\hat{y}_1' \cdot \hat{y}_2'). \tag{16}$$

7. Determine $\phi$, the angle between the roll–tilt axis and the mid–step $\hat{y}$ direction (the second column of $T$):

$$\phi = \cos^{-1}(\mathbf{rt} \cdot \hat{y}_{ms}). \tag{17}$$

8. Obtain roll $\rho$ and tilt $\tau$ via

$$\rho = \Gamma \cos(\phi), \tag{18}$$
$$\tau = \Gamma \sin(\phi). \tag{19}$$

9. Finally, determine shift, slide, and rise by projecting the positional difference onto the mid–step frame:

$$(D_x, D_y, D_z)^{\intercal} = T\,(\mathbf{r}_2 - \mathbf{r}_1)^{\intercal}. \tag{20}$$

#### DNA sequences

##### MNase-seq derived sequences

The MNase processed data were obtained from https://github.com/Jalbiti/NucleosomePeriodicity and analised using the pipeline developed by Orozco Lab [9]. Genome-wide nucleosome calls and nucleosome-depleted regions (NDRs) were obtained using nucleR package [10]. The genomic sequence corresponded to the S. cerevisiae S288C strain (R64-1-1 reference assembly).

Each nucleR nucleosome call carries a width score, indicating how well positioned the dyad is across cell populations, and a height score. Sequences with width and height scores $> 0.6$ and $> 0.4$, respectively, were classified as well-positioned nucleosomes. Each nucleosome-positioning sequence was defined as the midpoint of its NucleR call interval. The reported 211 bp sequence for each nucleosome spans dyad -105 to dyad +105. This window was chosen to place the nucleosome in the middle and include 32 bp of flanking DNA on each side for pulling simulations.

Candidate NDRs were taken from the -1 to +1 nucleosomes around the transcription start site (TSS). This restriction to promoter-proximal regions avoids unreliable gap-width estimates and poorly resolved nucleosomes.

Listed below are the DNA sequences used in this study. All sequences are reported in the 5'–3' direction.

**>1KX5**
GTTACATCCTGTGCATGTAAGTACTTACATGCACAGGATGTAACCTGCAGATACTACCAAAAGTGTA
TTTGGAAACTGCTCCATCAAAAGGCATGTTCAGCTGGATTCCAGCTGAACATGCCTTTTGATGGAGC
AGTTTCCAAATACACTTTTGGTAGTATCTGCAGGTGATTCTCCAGGGCGGCCAGTACTTACATGCAC
AGGATGTAAC

**>W601**
CCCTTATGTGATGGACCCTATACGCGGCCGCCCTGGAGAATCCCGGTGCCGAGGCCGCTCAATTGGT
CGTAGACAGCTCTAGCACCGCTTAAACGCACGTACGCGCTGTCCCCCGCGTTTTAACCGCCAAGGGG
ATTACTCCCTAGTCTCCAGGCACGTGTCAGATATATACATCCTGTGCATGTATTGAACAGCGACCTT
GCCGGTGCCA

**>ESRRB**
CACAGGGGACATTAATGGGTTATCAGCAGGGAGAAGGAGCGCCTCCCCATGTGGGACCTGGAGAAA
CAGAGGGTGGAGGGAGCATAGAGAGTCTGTTCTAAGCTGCAAAGCAAAGGCCTGGCGACCTAGGAG
ACCATGGAGTTCCAGAAAGTGATAGTTATGCAGAGCGAATGGAGGGAATCAGCACGCTGAGGCTGC
AGAGAGAGGGTGG

**>Lin28b**
TTTTACATCAGTTGGAGCATAAGTTAAGTGGTATTAACATATCCTCAGTGGTGAGTATTAACATGGA
ACTTACTCCAACAATACAGATGCTGAATAAATGTAGTCTAAGTGAAGGAAGAAGGAAAGGTGGGAG
CTGCCATCACTCAGAATTGTCCAGCAGGGATTGTGCAAGCTTGTGAATAAAGACACATACTTCATGT
AGTCAGAAGAG

>**Telomere**
TTAGGGTTAGGGTTAGGGTTAGGGTTAGGGTTAGGGTTAGGGTTAGGGTTAGGGTTAGGGTTAGGG
TTAGGGTTAGGGTTAGGGTTAGGGTTAGGGTTAGGGTTAGGGTTAGGGTTAGGGTTAGGGTTAGGG
TTAGGGTTAGGGTTAGGGTTAGGGTTAGGGTTAGGGTTAGGGTTAGGGTTAGGGTTAGGGTTAGGG
TTAGGGT

>**C2**
CTGGAGATACCCGGTGCTAAGGCCGCTTAATTCTGGAGATACCCGGTGCTAAGGCCGCTTAATTGGT
CGTAGCAAGCTCTAGCACCGCTTAAACGCACGTACGCGCTGTCTACCGCCTTTTAACCGCCAATAGG
ATTACTTACTAGTCTCTAGGCACGTGTAAGATATATACATGCTGTCTCTAGGCACGTGTAAGATATA
TACATCCTGT

>**5S-DNA**
GCCGGCCGGGCTGGGCTCTTGGGGCAGCCAGGCGCCTCCTTCAGCGTCTACGGCCATACCACCCTGA
ACGCGCCCGATCTCGTCTGATCTCGGAAGCTAAGCAGGGTCGGGCCTGGTTAGTACTTGGATGGGA
GACCGCCTGGGAATACCGGGTGCTGTAGGCTTTTTCTTTGGCTTTTTGCTGTTTCTTTCCTTTTCTT
CCAGACGGAGT

> $\alpha$**-satellite**
GCTTCCGTTTGCCTTTTATATGAACTTCCTGATATCAATATCCACCTGCAGATTCTACCAAAAGTGT
ATTTGGAAACTGCTCCATCAAAAGGCATGTTCAGCTCTGTGAGTGAAACTCCATCATCACAAAGAAT
ATTCTGAGAATGCTTCCGTTTGCCTTTTATATGAACTTCCTGATATCAATATCCACCTGCAGATTCT
ACCAAAAGTG

>**TP53 Promoter**
CAGCCCCAGCGATTTTCCCGAGCTGAAAATACACGGAGCCGAGAGCCCGTGACTCAGAGAGGACTCA
TCAAGTTCAGTCAGGAGCTTACCCAATCCAGGGAAGCGTGTCACCGTCGTGGAAAGCACGCTCCCAG
CCCGAACGCAAAGTGTCCCCGGAGCCCAGCAGCTACCTGCTCCCTGGACGGTGGCTCTAGACTTTTG
AGAAGCTCAA

>**PolyA**
AAAAAAAAAAAAAAAAAAAAAAAAAAAAAAAAAAAAAAAAAAAAAAAAAAAAAAAAAAAAAAAAAAA
AAAAAAAAAAAAAAAAAAAAAAAAAAAAAAAAAAAAAAAAAAAAAAAAAAAAAAAAAAAAAAAAAAA
AAAAAAAAAAAAAAAAAAAAAAAAAAAAAAAAAAAAAAAAAAAAAAAAAAAAAAAAAAAAAAAAAAA
AAAAAAAAAA

>**PolyAT**
ATATATATATATATATATATATATATATATATATATATATATATATATATATATATATATATATATAT
ATATATATATATATATATATATATATATATATATATATATATATATATATATATATATATATATATAT
ATATATATATATATATATATATATATATATATATATATATATATATATATATATATATATATATATAT
ATATATA

>**PolyTAA**
TAATAATAATAATAATAATAATAATAATAATAATAATAATAATAATAATAATAATAATAATAATAAT
AATAATAATAATAATAATAATAATAATAATAATAATAATAATAATAATAATAATAATAATAATAATA
ATAATAATAATAATAATAATAATAATAATAATAATAATAATAATAATAATAATAATAATAATAATAA
TAATAATAAT

>**PolyTAAA**
TAAATAAATAAATAAATAAATAAATAAATAAATAAATAAATAAATAAATAAATAAATAAATAAATAA
ATAAATAAATAAATAAATAAATAAATAAATAAATAAATAAATAAATAAATAAATAAATAAATAAATA
AATAAATAAATAAATAAATAAATAAATAAATAAATAAATAAATAAATAAATAAATAAATAAATAAAT
AAATAAATAA

>**PolyTAAAA**
TAAAATAAAATAAAATAAAATAAAATAAAATAAAATAAAATAAAATAAAATAAAATAAAATAAAATA
AAATAAAATAAAATAAAATAAAATAAAATAAAATAAAATAAAATAAAATAAAATAAAATAAAATAAA
ATAAAATAAAATAAAATAAAATAAAATAAAATAAAATAAAATAAAATAAAATAAAATAAAATAAAAT
AAAATAAAAT

>**PolyTAAAAA**
TAAAAATAAAAATAAAAATAAAAATAAAAATAAAAATAAAAATAAAAATAAAAATAAAAATAAAAAT
AAAAATAAAAATAAAAATAAAAATAAAAATAAAAATAAAAATAAAAATAAAAATAAAAATAAAAATA
AAAATAAAAATAAAAATAAAAATAAAAATAAAAATAAAAATAAAAATAAAAATAAAAATAAAAATAA
AAATAAAAAT

>**PolyTC**
TCTCTCTCTCTCTCTCTCTCTCTCTCTCTCTCTCTCTCTCTCTCTCTCTCTCTCTCTCTCTCTCTCTCT
CTCTCTCTCTCTCTCTCTCTCTCTCTCTCTCTCTCTCTCTCTCTCTCTCTCTCTCTCTCTCTCTCTCTC
TCTCTCTCTCTCTCTCTCTCTCTCTCTCTCTCTCTCTCTCTCTCTCTCTCTCTCTCTCTCTCTCTCTCT
CTCT
>**PolyTG**
TGTGTGTGTGTGTGTGTGTGTGTGTGTGTGTGTGTGTGTGTGTGTGTGTGTGTGTGTGTGTGTGTG
TGTGTGTGTGTGTGTGTGTGTGTGTGTGTGTGTGTGTGTGTGTGTGTGTGTGTGTGTGTGTGTGTG
TGTGTGTGTGTGTGTGTGTGTGTGTGTGTGTGTGTGTGTGTGTGTGTGTGTGTGTGTGTGTGTGTG
TGTGTGTGTGTGT
>**PolyG**
GGGGGGGGGGGGGGGGGGGGGGGGGGGGGGGGGGGGGGGGGGGGGGGGGGGGGGGGGGGGGGGGGG
GGGGGGGGGGGGGGGGGGGGGGGGGGGGGGGGGGGGGGGGGGGGGGGGGGGGGGGGGGGGGGGGGG
GGGGGGGGGGGGGGGGGGGGGGGGGGGGGGGGGGGGGGGGGGGGGGGGGGGGGGGGGGGGGGGGGG
GGGGGGGGGGGGG
>**PolyGC**
GCGCGCGCGCGCGCGCGCGCGCGCGCGCGCGCGCGCGCGCGCGCGCGCGCGCGCGCGCGCGCGCGC
GCGCGCGCGCGCGCGCGCGCGCGCGCGCGCGCGCGCGCGCGCGCGCGCGCGCGCGCGCGCGCGCGC
GCGCGCGCGCGCGCGCGCGCGCGCGCGCGCGCGCGCGCGCGCG
>**NDR; ACCESSION AB608241 REGION: 348..558**
TTTTTCTCTGATATCTGTAGCCCTAGTTTTTAATATTTTCGTAAAATGCTCAAAAAAATTTCTGTTTA
AATTCTTCTGGGGTATATACACATGTTCCCGTCAGCTAAATTAAAAAGAAGAAGGAACCGAAAGAAG
GCAAATTATAAAGTAATCATCCTCGGCATATTGAGGGGAAAATATCCGTCAAAAAAAAAAGTTCTGC
ACGTCGTCA
>**NPS; ACCESSION X71133 REGION: 27173..27383**
AAAAGAGCTTTCTATGTATGTATAGAGACAGCGTCTCAAAGGAAAAATTAGCCTCTTCGATGCCAAG
TACTTGTGATATACAACTGAAGAGGGCAATCAATGACGCTTATCCGGGCGGAGGAATAAAGGTCAC
GGTTCTGAACTCAACAACTGCAAGCTTGGACTCGTTAGCAACTACACATGTTAAGGAATTTGAGATT
GTTATCATTCC
>**Random; Low Elastic Deformation Energy**
GACAGGTACAAGAAGGAGTATGCATCAATGTGGTCGTGTGGAACAAACGCCACTGGAGACTGGGTT
AACCATTCGCTCCAGCGTCATGAAAGTCACTGTTAGGGCGACCTTCGATTCGGATGTGACATTTCAT
TACATTACGCTCAGGACTGCGAACGAAAGATTTAAGAATGCTTAACCCGGTACCTAACCCATCTGAT
TTTTACACACATC
>**Random; Intermediate Elastic Deformation Energy**
AGTAGCTCTGGCACGTGCCGGGGACAGCGCTTCAGTGCCTTTGGGTTCTAGCGCCGCGATATCTATA
CACATCCGGCGTGTTGCCATCATAATATCCGAAGTCTTAACTAGAGGACCACAAGCCGATCTGAAGC
GGTAACCGGGTGTATCTTCAGAGTCATTCAGGATTAGTTTCCCGTCTAGGCCGCTACCTGCATGAGG
AGGATACGAC
>**Random; High Elastic Deformation Energy**
GGTTACTATCAACCACGTCCACCATCGTCCAAGGACGAACGAAACCTTCTTCCAGAAGATTCCGGCA
CACTCGTTTGAAGGAAAGTTCTCGGTTCCCCACGTTCAAAACCTAGGCCGGAGAATTACTTCAACGT
TCGTCTCAGTCCTGACTGCAGAACGGATTATCTCCACAGAAAGGCACAGACCGGGCCCCTCACACGT
ACTGCGAGCC

#### Supplementary Tables

**Supplementary Table 1.** Experimental and simulation force measurements for nucleosome unwrapping under applied force. F1 and F2 denote the first and second force maxima corresponding to outer- and inner-turn unwrapping, respectively.

| Reference | F1 (pN) | F2 (pN) | Conditions / Notes |
|---|---|---|---|
| Simulation (this work) | $6.14 \pm 0.48$ | $27.87 \pm 2.17$ | Widom 601 sequence; 150 mM monovalent salt |
| Díaz-Celis *et al.*, 2022 [11] | $3.9 \pm 0.3$ | $30.4 \pm 8.3$ | 50 mM NaCl; Widom 601; mononucleosome |
| Díaz-Celis *et al.*, 2022 [11] | $3.2 \pm 0.4$ | $24.4 \pm 4.2$ | 200 mM NaCl; Widom 601; mononucleosome |
| Ngo *et al.*, 2015 [12] | 3–5 | $^{\dagger}16.8 \pm 1.5$ | 50 mM NaCl, 1 mM $MgCl_2$; Widom 601; mononucleosome |
| Chien *et al.*, 2014 [13] | 2–3.5 | — | 10–200 mM $K^+$; chromatin fiber |
| Mihardja *et al.*, 2006 [14] | ∼3 | — | 50 mM KOAc, 10 mM magnesium acetate, 1 mM DTT, 0.1 mg/ml BSA; Widom 601; chicken erythrocyte histones |
| Brower-Toland *et al.*, 2002 [15] | — | 17.6 | 100 mM NaCl, 10 mM Tris-HCl (pH 8.0), 1 mM $Na_2EDTA$, 1.5 mM $MgCl_2$, 0.02% (v/v) Tween 20, 0.01% (w/v) milk protein, 22°C; 5S rDNA; nucleosome arrays |
| Brower-Toland *et al.*, 2002 [15] | — | 20.2 | Same buffer conditions as above; 5S rDNA; nucleosome arrays |

$^{\dagger}$F2 value corresponds to one-end release only.

**Supplementary Table 2.** Unwrapping free energies ($\Delta\Delta G$), standard error of the mean (s.e.m), and absolute difference relative to the canonical nucleosome (1KX5).

| Condition | $\Delta\Delta G$ (kcal/mol) | s.e.m | $\Delta\Delta G^{\mathrm{mod}} - \Delta\Delta G^{\mathrm{1KX5}}$ (kcal/mol) |
|---|---|---|---|
| 1KX5 (Canonical) | 63.49 | 1.81 | — |
| Hyperacetylation (Core + Tails) | 11.68 | 1.50 | $-51.81 \pm 2.35$ |
| Hyperacetylation (Core) | 28.64 | 1.91 | $-34.85 \pm 2.63$ |
| Hyperacetylation (Tails) | 30.56 | 3.52 | $-32.93 \pm 3.96$ |
| Hyperacetylation (Endogenous) | 52.52 | 1.47 | $-10.97 \pm 2.34$ |
| Hyperacetylation (H3 + H2AC tails) | 48.10 | 3.15 | $-15.39 \pm 3.64$ |
| H4K16Ac | 57.96 | 1.23 | $-5.53 \pm 2.18$ |
| H3K56ac | 59.59 | 1.61 | $-3.90 \pm 2.42$ |
| H3K56ac/H4K77ac/K79ac | 50.63 | 1.07 | $-12.86 \pm 2.10$ |
| H3.3 | 66.02 | 1.76 | $+2.53 \pm 2.53$ |
| CENP-A | 59.83 | 2.43 | $-3.66 \pm 3.02$ |
| H2A.Z | 56.79 | 1.01 | $-6.70 \pm 2.08$ |
| macroH2A | 70.47 | 2.52 | $+6.98 \pm 3.10$ |

**Supplementary Table 3.** Peak forces associated with the mechanical barriers for outer and inner turn unwrapping for nucleosomes with modified DNA sequences ($F_1^{\mathrm{mod}}$, $F_2^{\mathrm{mod}}$), percentage differences ($\Delta\%F_1^{\mathrm{mod}}$, $\Delta\%F_2^{\mathrm{mod}}$) relative to the canonical nucleosome ($F_1^{\mathrm{1KX5}}$, $F_2^{\mathrm{1KX5}}$), and the change in extension at the second mechanical barrier ($\Delta X_{\mathrm{F2}}$).

| System | $F_1^{\mathrm{mod}}$ (pN) | $F_2^{\mathrm{mod}}$ (pN) | $\Delta\%F_1^{\mathrm{mod}}$ | $\Delta\%F_2^{\mathrm{mod}}$ | $\Delta X_{\mathrm{F2}}$ (Å) |
|---|---|---|---|---|---|
| 1KX5 (canonical) | $6.95 \pm 0.36$ | $35.77 \pm 0.92$ | — | — | — |
| W601 | $6.14 \pm 0.48$ | $27.87 \pm 2.17$ | $-11.6 \pm 8.3$ | $-22.1 \pm 6.4$ | $0.00 \pm 4.48$ |
| C2 | $6.53 \pm 0.47$ | $28.36 \pm 0.72$ | $-6.1 \pm 8.4$ | $-20.7 \pm 2.9$ | $-10.43 \pm 2.36$ |
| polyTC | $4.92 \pm 0.45$ | $32.37 \pm 2.34$ | $-29.1 \pm 7.5$ | $-9.5 \pm 6.9$ | $+24.32 \pm 6.35$ |
| polyG | $4.95 \pm 0.18$ | $25.12 \pm 1.15$ | $-28.8 \pm 4.5$ | $-29.8 \pm 3.7$ | $+17.38 \pm 1.30$ |
| polyA | $5.82 \pm 0.62$ | $31.17 \pm 1.90$ | $-16.2 \pm 10.0$ | $-12.9 \pm 5.8$ | $+10.42 \pm 1.70$ |
| polyGC | $6.54 \pm 0.14$ | $29.39 \pm 1.74$ | $-5.8 \pm 5.3$ | $-17.8 \pm 5.3$ | $-3.48 \pm 2.46$ |
| polyTG | $7.01 \pm 0.21$ | $25.91 \pm 1.44$ | $+0.9 \pm 6.0$ | $-27.6 \pm 4.4$ | $-38.23 \pm 0.85$ |
| polyAT | $7.16 \pm 0.43$ | $25.48 \pm 0.45$ | $+3.1 \pm 8.2$ | $-28.8 \pm 2.2$ | $-41.70 \pm 3.18$ |
| 5S-DNA | $7.44 \pm 0.53$ | $29.22 \pm 1.61$ | $+7.1 \pm 9.4$ | $-18.3 \pm 5.0$ | $+10.42 \pm 10.51$ |
| Telomere | $6.19 \pm 0.62$ | $27.66 \pm 0.99$ | $-10.8 \pm 10.1$ | $-22.7 \pm 3.4$ | $+3.47 \pm 2.36$ |
| Lin28b | $6.36 \pm 0.37$ | $29.84 \pm 1.56$ | $-8.4 \pm 7.1$ | $-16.6 \pm 4.9$ | $-6.95 \pm 10.51$ |
| ESRRB | $6.75 \pm 0.46$ | $27.02 \pm 0.29$ | $-2.8 \pm 8.4$ | $-24.5 \pm 2.1$ | $-6.95 \pm 10.17$ |

**Supplementary Table 4.** Peak forces associated with the mechanical barriers for outer and inner turn unwrapping for nucleosomes incorporating different histone modifications and DNA sequences ($F_1^{\mathrm{mod}}$, $F_2^{\mathrm{mod}}$), and percentage differences ($\Delta\%F_1^{\mathrm{mod}}$, $\Delta\%F_2^{\mathrm{mod}}$) relative to the canonical nucleosome ($F_1^{\mathrm{1KX5}}$, $F_2^{\mathrm{1KX5}}$).

| System | $F_1^{\mathrm{mod}}$ (pN) | $F_2^{\mathrm{mod}}$ (pN) | $\Delta\%F_1^{\mathrm{mod}}$ | $\Delta\%F_2^{\mathrm{mod}}$ |
|---|---|---|---|---|
| 1KX5 (canonical) | $6.95 \pm 0.36$ | $35.77 \pm 0.92$ | — | — |
| 1KX5 + alpha | $6.93 \pm 0.34$ | $26.72 \pm 1.02$ | $-0.3 \pm 7.1$ | $-25.3 \pm 3.4$ |
| CENP-A | $2.94 \pm 0.04$ | $36.92 \pm 2.65$ | $-57.7 \pm 2.3$ | $+3.2 \pm 7.9$ |
| CENP-A + alpha | $4.25 \pm 1.00$ | $35.72 \pm 1.52$ | $-38.8 \pm 14.8$ | $-0.1 \pm 5.0$ |
| 1KX5 + TP53 | $5.54 \pm 0.41$ | $28.13 \pm 0.91$ | $-20.3 \pm 7.2$ | $-21.4 \pm 3.2$ |
| H2A.Z | $9.46 \pm 0.53$ | $28.58 \pm 0.65$ | $+36.2 \pm 10.4$ | $-20.1 \pm 2.7$ |
| H2A.Z + TP53 | $5.76 \pm 0.62$ | $24.81 \pm 1.14$ | $-17.1 \pm 9.9$ | $-30.6 \pm 3.6$ |
| Hyperacetylation (Core + Tails) | $2.21 \pm 0.53$ | $5.83 \pm 0.41$ | $-68.2 \pm 7.8$ | $-83.7 \pm 1.2$ |
| Hyperacetylation (Core) | $2.15 \pm 0.42$ | $13.02 \pm 0.96$ | $-69.1 \pm 6.2$ | $-63.6 \pm 2.8$ |
| Hyperacetylation (Tails) | $2.37 \pm 1.07$ | $18.88 \pm 2.33$ | $-65.8 \pm 15.5$ | $-47.2 \pm 6.6$ |
| Hyperacetylation (Endogenous) | $8.48 \pm 0.77$ | $26.99 \pm 0.82$ | $+22.0 \pm 12.8$ | $-24.5 \pm 3.0$ |
| Hyperacetylation (H3 + H2AC tails only) | $5.14 \pm 0.34$ | $28.69 \pm 2.19$ | $-26.0 \pm 6.2$ | $-19.8 \pm 6.5$ |
| H3K56ac | $5.27 \pm 0.73$ | $33.97 \pm 2.95$ | $-24.1 \pm 11.2$ | $-5.0 \pm 8.6$ |
| H3K56ac/H4K77ac/K79ac | $5.60 \pm 0.41$ | $21.91 \pm 0.73$ | $-19.4 \pm 7.3$ | $-38.7 \pm 2.6$ |
| H4K16Ac | $6.48 \pm 0.42$ | $35.31 \pm 1.98$ | $-6.7 \pm 7.8$ | $-1.3 \pm 6.1$ |
| polyA | $5.82 \pm 0.62$ | $31.17 \pm 1.90$ | $-16.2 \pm 10.0$ | $-12.9 \pm 5.8$ |
| H3 + H2AC tails acetylation + polyA | $3.65 \pm 0.26$ | $32.17 \pm 1.95$ | $-47.5 \pm 4.7$ | $-10.1 \pm 5.9$ |
| polyAT | $7.16 \pm 0.43$ | $25.48 \pm 0.45$ | $+3.1 \pm 8.2$ | $-28.8 \pm 2.2$ |
| H3 + H2AC tails acetylation + polyAT | $5.90 \pm 0.14$ | $24.68 \pm 1.41$ | $-15.0 \pm 4.8$ | $-31.0 \pm 4.3$ |
| H3.3 | $7.66 \pm 0.36$ | $36.12 \pm 1.64$ | $+10.3 \pm 7.8$ | $+1.0 \pm 5.3$ |
| macroH2A | $11.99 \pm 0.65$ | $30.70 \pm 2.98$ | $+72.6 \pm 13.0$ | $-14.2 \pm 8.6$ |

**Supplementary Table 5.** For all conditions, the minimum overlap between adjacent windows exceeds 0.1, ensuring sufficient histogram overlap for reliable WHAM reconstruction. The last column reports the aggregated simulation time per condition.

| Condition | $n$ windows | min overlap | mean overlap | Total time ($\mu$s) |
|---|---|---|---|---|
| 1KX5 | 300 | 0.244 | 0.821 | 12 |
| 1KX5 (additional 100 replicates) | 6000 | 0.322 | 0.9576 | 240 |
| Hyperac. (Core + Tails) | 300 | 0.240 | 0.826 | 12 |
| Hyperac. (Core) | 300 | 0.243 | 0.822 | 12 |
| Hyperac. (Tails) | 300 | 0.243 | 0.823 | 12 |
| Hyperac. (tails endogenous) | 300 | 0.231 | 0.820 | 12 |
| Hyperac. (H3 + H2AC tails) | 300 | 0.204 | 0.823 | 12 |
| H4K16ac | 300 | 0.115 | 0.822 | 12 |
| H3K56ac | 300 | 0.177 | 0.813 | 12 |
| H3K56ac + H4K77ac + K79ac | 300 | 0.236 | 0.822 | 12 |
| CENP-A | 300 | 0.157 | 0.822 | 12 |
| H2A.Z | 300 | 0.240 | 0.822 | 12 |
| H3.3 | 300 | 0.132 | 0.819 | 12 |
| macroH2A | 300 | 0.238 | 0.817 | 12 |
| KXA | 300 | 0.233 | 0.822 | 12 |
| KXQ | 300 | 0.242 | 0.826 | 12 |
| RXA | 300 | 0.218 | 0.824 | 12 |
| RXQ | 300 | 0.215 | 0.829 | 12 |
| TAA | 300 | 0.214 | 0.821 | 12 |
| TAAA | 300 | 0.151 | 0.823 | 12 |
| TAAAA | 300 | 0.231 | 0.821 | 12 |
| TAAAAA | 300 | 0.236 | 0.820 | 12 |
| polyAT | 300 | 0.236 | 0.826 | 12 |
| polyA | 300 | 0.244 | 0.819 | 12 |
| polyG | 300 | 0.239 | 0.821 | 12 |
| polyGC | 300 | 0.233 | 0.824 | 12 |
| polyTG | 300 | 0.238 | 0.826 | 12 |
| polyTC | 300 | 0.227 | 0.821 | 12 |
| C2 | 300 | 0.238 | 0.821 | 12 |
| ESRRB | 300 | 0.232 | 0.820 | 12 |
| lin28b | 300 | 0.239 | 0.822 | 12 |
| W601 | 300 | 0.236 | 0.822 | 12 |
| 5S-DNA | 300 | 0.229 | 0.822 | 12 |
| Telomere | 300 | 0.237 | 0.822 | 12 |
| NDR | 300 | 0.332 | 0.922 | 12 |
| NPS | 300 | 0.448 | 0.920 | 12 |
| Random; Low E.D. Energy | 300 | 0.227 | 0.923 | 12 |
| Random; Int. E.D. Energy | 300 | 0.632 | 0.925 | 12 |
| Random; High E.D. Energy | 300 | 0.292 | 0.925 | 12 |
| 1KX5 + alpha | 300 | 0.236 | 0.823 | 12 |
| CENP-A + alpha | 300 | 0.230 | 0.820 | 12 |
| 1KX5 + TP53 | 300 | 0.234 | 0.822 | 12 |
| H2A.Z + TP53 | 300 | 0.233 | 0.819 | 12 |
| polyAT + Hyperac. (H3 + H2AC tails) | 300 | 0.241 | 0.824 | 12 |
| polyA + Hyperac. (H3 + H2AC tails) | 300 | 0.248 | 0.818 | 12 |
